# Partner cues and individual variation underlie sex-reversed parental care in poison frogs

**DOI:** 10.1101/2025.09.02.672740

**Authors:** Jeanette B. Moss, Brittany M. Winter, Sarah E. Westrick, Katie Julkowski, Molly E. Podraza, Eva K. Fischer

## Abstract

Flexible parental care strategies are widespread in nature and factor into conflict between the sexes and the realization of sex roles. While adaptive explanations abound, the mechanisms underlying flexible ‘sex-reversal’ of care are less clear. We enlist a biparental frog (*Ranitomeya imitator*) with flexible parental care to investigate the extrinsic and intrinsic mechanisms underpinning parental decisions. Using mate removal experiments in the laboratory, we show that members of the primary caregiving sex (males) show less variation than the flexible sex (females) in their propensity to provide care, and that care propensity in females is affected by extrinsic partner cues as well as individual variability. Indeed, individual repeatability in parental effort is high in both typically caregiving and flexible parents. To investigate the underpinnings of differences in care propensity, we sequenced RNA from whole brains of caregiving and non-caregiving frogs of both sexes. While actively caregiving females showed minimal differential gene expression compared to actively caregiving males, females that failed to provide care showed distinct patterns of gene expression. Our findings offer an initial glimpse into the environmental and genetic regulation of individual variation in sex-reversed parental care.

## Introduction

Cross-sexual transfer, a process by which traits ancestrally expressed in only one sex appear in the opposite sex in descendant lineages [1], provides a powerful framework for studying the evolution of behavior, including parental care [2,3]. While parenting systems are often characterized by sex-specific behaviors (i.e., ‘sex roles’) that can be remarkably stable over evolutionary timescales [4–8], there is also inherent flexibility required for rearing offspring [9]. Selection may capitalize on the mechanisms mediating behavioral flexibility within species to drive the evolution of different care strategies between species via cross-sexual transfer [2,3]. While studies across taxa suggest that males and females rely on largely shared molecular ‘toolkits’ for care [10–13], the conditions that lead to their activation may differ. Thus, understanding the conditions that promote evolutionary reversal of parental sex roles requires probing the mechanisms underlying both their maintenance and plasticity in ancestral lineages. Given that the capacity for sex-reversed parental care is not ubiquitous and appears to vary among individuals, contexts, and species, investigations that incorporate individual behavioral variation will provide new insights into the extrinsic factors and underlying mechanisms that promote cross-sexual transfer of parental care.

Neotropical poison frogs (family: Dendrobatidae) offer an exciting system for investigating cross-sexual transfer of parental behavior owing to diversity of care modes among closely related species combined with behavioral flexibility within species [3,14–17]. While there is tremendous diversity in *who* expresses specific behaviors, care follows a similar sequence across many species: (1) parents care for and guard terrestrial eggs; (2) hatched tadpoles are transported “piggyback” style to water to complete metamorphosis; and in some species, (3) mothers return to lay unfertilized, trophic eggs for tadpoles to consume (reviewed in [18]). Based on ancestral state reconstructions in this group, West-Eberhard (2003) and others [19,20] have theorized that uniparental male care likely gave rise to uniparental female care *via* biparental care with female behavioral flexibility as a stepping stone. If so, then sex-specific plasticity in parental behavior lays the foundation for cross-sexual transfer of care. Studies of the extrinsic and intrinsic factors driving parental compensation in biparental species should offer clues to the evolutionary substrates that facilitate transitions in care mode.

As in many biparental species, biparental poison frogs subdivide care tasks, with one sex serving as the primary caregiver [21–25]. In the monogamous thumbnail dart frog, *Ranitomeya imitator*, males and females share the duty of egg care [25] and females facultatively provision tadpoles with trophic eggs [24,26]. Although mothers associate closely with male partners throughout care and return to deposition pools to feed tadpoles, the act of tadpole transport is performed overwhelmingly by males, who also coordinate egg feeding by alerting females to offspring need through specialized calls [22,24,27]. Yet recent empirical work and our own observations confirm a capacity for flexible sex-reversal in this and related species [12,24,28]. Specifically, females may be induced to takeover tadpole transport when their partner is removed. Based on field observations of *Allobates femoralis*, Ringler *et al*. (2015) proposed that male acoustic signals may play a key role in informing female takeover decisions. In *R. imitator*, males produce distinct calls to advertise and defend territories, court females, and solicit trophic egg feeding [27,29,30] and there is some evidence that reliance on acoustic communication for effective caregiving is associated with the evolution of individual vocal recognition and partner preference [31]. Nevertheless, the extrinsic cues that facilitate recognition and communication among parents are likely multimodal [27].

While the propensity for parental care differs between the sexes, underlying mechanisms may be shared in males and females [10–13]. Recent investigations into the hormonal and neurogenomic basis of care in poison frogs have revealed similar patterns of glucocorticoid response and neural activity in transporting males and females [12,32]; however, the mechanisms mediating behavioral plasticity in the non-typically transporting sex remain unclear. Interestingly, work in the uniparental male species, *Dendrobates tinctorius*, has shown that observing female parents exhibit hormone and brain transcriptome – but not neural activity – profiles paralleling those of their actively transporting male partners [12]. One possible explanation for this pattern is that, in addition to ‘priming’ for possible takeover of care duties through activation of genes associated with transport, observing parents recruit additional gene sets to monitor the progression of care directed by their partner and make decisions as to whether to takeover. In biparental *R. imitator*, non-transporting females even exhibit similar patterns of neural induction to their actively transporting partners, presumably reflective of an overall ‘caring’ state in females that provide egg care and perform egg feeding even though they do not typically transport tadpoles [32].

We investigated the mechanisms of sex-reversed tadpole transport in biparental *R. imitator*. We began by carrying out mate removal experiments to characterize sex and individual variation in transport behavior and determine the relative contributions of male visual and acoustic signals to female takeover decisions. We predicted that males and females would differ in their propensity for and performance of a male-biased parental task (tadpole transport) but not shared tasks (egg care), and that individual variation in propensity to transport would be inflated in the typically non-transporting sex (females). Further, we anticipated that partner social cues (male presence and isolated acoustic and visual signals) would impose strong effects on female transport decisions, but that male transport would not differ in the presence vs absence of females. To explore underlying mechanisms associated with behavioral differences, we then compared hormones and brain gene expression profiles of males and females that flexibly provide care to females that did not. We expected that both sexes would show changes in glucocorticoid circulation and gene expression associated with the activation of transport behavior, and that variation between observing, transporting, and non-transporting females would pinpoint mechanisms underlying the capacity for sex-reversed care.

## Methods

## (a) Frog Husbandry

Frogs for this study were from a captive colony maintained at the University of Illinois Urbana-Champaign (UIUC). Tanks (12×12×18 inch glass terraria with mesh lids (©Exo Terra, Mansfield, MA, U.S.A)) were located in temperature-(22.05 ± 2.47°C) and light-(12:12 h cycle) controlled rooms and ambient humidity (84.95 ± 3.08%) was maintained via a misting system (©Mist King, Ontario, Canada). Terraria were outfitted with live plants, driftwood, sphagnum moss, leaf litter, and film canisters: two horizontal canisters positioned ∼10” above the ground for egg laying and two vertical, water-filled canisters on the ground for tadpole deposition. In captivity as in the wild, once males and females have established a pair bond, females lay clutches of 1–4 terrestrial eggs as frequently as every 1–2 weeks (Brown *et al*., 2008, pers. obs.). Adults were housed as breeding pairs and monitored daily for signs of breeding activity (e.g., eggs and/or tadpoles), which were documented in a digital database. Additionally, tanks were continuously monitored using a video surveillance system (RLC-510A, ©Reolink, New Castle, DE, U.S.A.), with videos backed up manually to a server. Frogs were fed flightless *Drosophila* fruit flies dusted with vitamin supplements three times weekly. All procedures were approved by the UIUC Animal Care and Use Committee (Protocol #20147).

### (b) Behavior

#### i. Experimental Overview

To investigate the role of social stimuli in mediating sex-reversed tadpole transport, we conducted six types of trials. The first trial served as a behavioral baseline (Control), wherein male and female behavior was observed in the absence of any manipulation. In subsequent trials, either the male or the female was removed to observe the behavior of single parents. To determine whether a female’s propensity to takeover tadpole transport is affected by cues of male presence (visual and/or auditory), we also conducted trials in which males were removed and females were presented with a 3D-printed male dummy painted to match their mate’s coloration and pattern (Appendix A; Fig. S1), playback of their mate’s advertisement calls (Appendix B), or both. Only pairs that had successfully produced eggs and transported tadpoles on at least two occasions were used in this experiment. Trial order was randomized following the Control (baseline) trial. Additional life-history details for focal individuals are in Table S1.

### ii. Data Collection and Monitoring

During the trial period from October 2021 through June 2023, *R. imitator* breeding activity was monitored to identify pairs and clutches suitable for trials. Trials were initiated on the eighth day (day 8) following the discovery of a clutch (day 0) in the home terrarium. Focal pairs always began with a Control trial, and once a behavioral baseline was established, entry into the other four trial types was randomized. For mate removal trials (male removal and female removal), either the male or the female was moved to temporary housing in a non-adjoining room to prevent any acoustic communication between partners during the trial. For visual only trials, the focal male was removed and replaced with his corresponding dummy (Appendix A), which was affixed to the glass wall of the terrarium immediately adjacent to the egg canister. For audio only trials, a speaker (Mod1 Orb speaker, Orb Audio, New York, NY, USA) was suspended with its woofer facing down approximately two inches above the mesh lid using a cutout wooden frame. Recorded calls belonging to the focal male (Appendix B) were stored on individual USB drives such that at the end of each 2-hour sequence, the track restarted from the beginning and looped for the duration of the trial. Calls were broadcast through an amplifier (MAMP1, MouKey, Solihull, UK) at 60–70 dB, which approximates the sound pressure level (SPL) of advertising males measured from a distance of ∼30 cm. The frequency response of the playback system was flat ±2.5 dB over the range of interest (2–6 kHz). Acoustic signals were only broadcast during daytime hours when diurnal *R. imitator* are active (0600–1800). To initiate visual + auditory trials, dummies and playback were presented as described above but simultaneously. Video/audio recording of daylight hours (0600–1800) began on day 9.

From day 9 onwards, focal clutches were inspected once daily for signs of hatching. Hatch dates were recorded as the day on which the first hatched tadpole was detected (on average, 12.9 ± 2.6 days post-oviposition). Failure to hatch (i.e., due to molding or desiccation) resulted in termination of the trial and eventual retrial. From the time of hatching, egg canisters and deposition pools were checked daily for evidence of transport and tanks were visually inspected for transporting frogs. We avoided manually searching tanks so as not to stress the frogs and disrupt normal parental behaviors. Based on our preliminary data, we concluded that hatched tadpoles typically desiccate within approximately four days if not transported. Therefore, if no transport occurred within five days (excluding hatch day), the trial was ended, and we recorded the outcome as failure to transport. Transport activity was recorded from the time a hatched tadpole disappeared from the egg canister, was visually observed on a frog, and/or appeared in a pool. The disappearance of all hatched tadpoles paired with their failure to appear on a parent or in a pool within three days prompted us to end the trial and record the outcome as a failure to transport. If frogs were detected with tadpoles on their backs, they were given as many days as needed to deposit the tadpole in a pool (successful transport) or until they lost the tadpole (failure to transport). As soon as the first transported tadpole was detected in a pool, we noted the time and location and ended the trial. All stimuli were removed from the tank, video recording stopped, and mates returned. After pairs were reunited, subsequent trials were only conducted on newly laid clutches, such that all pairs were afforded at least eight days between the completion of one trial and the initiation of another.

#### iii. Video scoring

At a trial’s conclusion all recorded videos were reviewed to confirm the identity of the transporter (in the case of Control trials) and refine the time window for transport. For trials involving two focal individuals (i.e., male and female pair), individuals were differentiated based on their distinct color patterns. To determine whether prehatching parental behaviors predict transport, we screened video from the last full 12-hr light cycle before hatch day. Because some clutches hatched as early as day 9 or 10 and recording began on day 9, this was the best way to standardize the timing of scoring across clutches while maximizing the number of scorable trials. For each focal individual, we quantified the number of times they visited the focal egg canister (defined as full body inside of the canister) and the duration of each visit (in minutes), as well as the number and duration of visits to deposition pools (defined as full body inside the pool).

From hatch day onwards, videos were screened for evidence of tadpole transport. Because the angle of the cameras made it impossible to see inside of egg canisters, we recorded the timing of tadpole pickup as the last time the focal frog entered the egg canister before it emerged with a tadpole on its back. Transporting frogs were observed until they entered a pool and exited without a tadpole or otherwise appeared in frame without a tadpole on their back, whichever came first. Timing of deposition was recorded as the last time the focal frog entered a deposition pool before emerging without a tadpole. We also recorded the number of visits to pools (i.e., attempted depositions) before successful deposition. Trials in which frogs failed to transport based on direct inspection of tanks were reviewed for evidence of tadpole pickup.

#### iv. Audio scoring

We quantified male call rates across three distinct periods of parental care in the presence and absence of female partners. We predicted that if calling functions primarily within pair bonds to communicate changes in offspring state, then males should reduce both the effort they expend on calling (i.e., overall call rate) and degree to which they modulate calling (i.e., variation in call rate across care) when females are removed. To test this, we selected a subset of ten trials (N=5 control and N=5 female removed) with known hatch dates and transport times and for which the same five males were repeated to account for individual variation in calling behavior. We did not score audio tracks for every trial as the scrutiny required for accurate call enumeration rendered this prohibitive at scale (Appendix C). We extracted audio from video recordings corresponding to three time periods: (1) the last full (12-hr) day before hatch day, or *pre-hatching*; (2) the day of hatching up to the moment of tadpole pickup as identified from the video analysis, or *pre-transport*; (3) the time that the tadpole was in transit (i.e., pick-up to drop-off) as identified from the video analysis, or *transport*.

Scoring was conducted blind to treatment groups and sampling periods. We made no distinctions between call ‘types’ due to strong overlap in acoustic properties [27]. Call rate was quantified by dividing the total number of calls by the duration of the trimmed and filtered track (in hours).

#### v. Data Analysis

All statistical analyses were performed in R v. 4.4.2 (http://www.R-project.org/). To contrast parenting behaviors between the sexes and across trial types, we fit a series of linear mixed models (lmm) and generalized linear mixed models (glmm) with the package, ‘lme4’ [33].

Male call rates during trials were modeled as a Poisson regression with treatment (female present and female removed), parental stage (pre-hatching, hatch day, and transport), and their interaction as fixed effects and male ID as a random effect. We compared transport success between treatments using binomial regression models, in which transporting sex and trial type were specified as fixed effects and transporter ID as a random effect. To test for order effects, an additional model was constructed including the chronological sequence of the trial (e.g., for an individual transporter, an integer between 1 and 4) as a covariate. To specifically address whether females became more likely to takeover transport with experience, we fit a model of transport success in trials 2–4 in which transport success in trial 1 (the first mate removal trial) was specified as a fixed effect. Pairwise post hoc testing of each model was carried out using the ‘emmeans’ package with Tukey adjustment [34]. Individual repeatability in transport success, adjusted for the fixed effect of trial type, was assessed using the rptBinary function in the R package, ‘rptR’ with 1000 parametric bootstraps [35]. To evaluate sex differences in the latency to transport tadpoles, we fit Cox proportional hazard regression models as implemented in the package ‘survival’ [36]. Tadpole transit time (i.e., from tadpole pickup to tadpole drop-off) was modeled as a Gamma regression with transporting sex as a fixed effect and transporter ID as a random effect.

To investigate variation in prehatching behaviors, we inspected the distribution of egg and pool visitation rates (0–16) and their cumulative duration (0–12,240 s) in the last full day before egg hatching. The duration of egg and pool visits was log-transformed to meet assumptions of normality, and individual repeatability in each behavior was assessed using the rptPoisson or rptGaussian functions in ‘rptR,’ as appropriate. We modeled variation in the number of visits to eggs and pools and the log-transformed duration of egg and pool visits using glmms with Poisson links and lmms, respectively. Effects of sex and trial type on each behavior were assessed by specifying these variables as fixed effect in the models, with the latter being carried out separately for each sex. We next tested whether any prehatching behaviors would predict transport propensity among females by fitting a series of binomial regressions with the behavior as a fixed effect. Finally, we examined the effect of prehatching behavior on latency to transport and log-transformed tadpole transit time among frogs that successfully transported tadpoles. Latency to transport and transit time were modeled using glmms with Poisson links and lmms, respectively, and prehatching behavior, sex, and their interaction were included as fixed effects. In all models, individual ID was included as a random effect.

### (c) Hormone and Gene Expression Analyses

#### i. Tissue Collection

To investigate the hormonal and molecular mechanisms associated with flexible transport across the sexes, a subset of frogs (N=21 females and N=10 males) was selected for whole brain gene expression analysis. Due to mortality of some individuals used for behavior, additional individuals were assayed (i.e., transport propensity was recorded during a mate removal trial) for brain gene expression. Based on observed individual repeatability in female transport propensity, females were categorized *a priori* as either Flexible (i.e., transported tadpole in a male removal trial) or Inflexible (i.e., failed to transport tadpole in a male removal trial). For terminal sampling trials, focal clutches were monitored each day as in behavioral trials and male partners were removed on Day 9 to create the Sex-Reversed social condition.

Frogs were sacrificed at three timepoints: (1) during egg care, but before hatching; (2) after eggs hatched and tadpole transport was initiated (within one day of hatching); and (3) after tadpoles had been hatched for three days without transport. Based on data from behavioral trials, 82% of tadpole transport events occurs within two days of eggs hatching. Thus, three days ensured that tadpoles were still alive and available for transport (i.e., offspring cues still present) but the likelihood that frogs were sacrificed before they had the opportunity to transport was low. This sampling scheme yielded four behavioral groups: Egg Care (i.e., males (N=5) and females (N=5) performing egg care only, sampled 9 days post-oviposition), Sex-Typical (i.e., male transporting (N=5) and female observing (N=5), sampled 0–1 days post-hatching), Sex-Reversed-Flexible (i.e., female transporting in absence of male (N=5), sampled 0–1 days post-hatching), and Sex-Reversed-Inflexible (i.e., non-transporting female in absence of male (N=5), sampled 3 days post-hatching). Because hatching times varied between clutches, timing of sampling varied slightly across behavioral groups. Relative to oviposition day (Day 0), sex-typical pairs were sampled between day 10 and day 15 and sex-reversed females were sampled between day 14 and day 20 (Flexible females: day 14–20; Inflexible females: day 15– 17).

Immediately upon capture, frogs were rapidly decapitated, and trunk blood siphoned with heparinized capillary tubes. Blood was centrifuged at 4°C to separate plasma, which was stored frozen at -20°C until further processing. Due to *R. imitator*’s small body size, plasma collection was not possible in all cases. Whole brains were dissected simultaneously with blood collection. Tissue was added to cryotubes pre-filled with ceramic beads (Omni International, Kennesaw, GA, USA) and flash-frozen in liquid nitrogen. Brain tissue was stored at -80°C until further processing. This full process took <5 minutes.

#### ii. Hormone Quantification and Analysis

Plasma samples were assayed for cortisol using a competitive ELISA immunoassay kit (K003-H; Arbor Assays, Ann Arbor, MI, U.S.A). We chose cortisol because our research group recently demonstrated that cortisol is the more prevalent glucocorticoid present in waterborne hormone samples for *R. imitator* [37] and is associated with parental behavior in *D. tinctorius* [12]. Each sample was run in duplicate by adding equal volume of dissociation reagent to the plasma, then letting it sit at room temperature for five minutes before resuspension in assay buffer. Plasma volume used in the assay differed due to variation in the volume of plasma collected. Samples with 1 uL of plasma (n=2), 2 uL of plasma (n=7), 4 uL of plasma (n=1), and 5 uL of plasma (n=17) were resuspended in 198, 196, 392, or 390 uL of assay buffer, respectively. Assays were carried out following manufacturer instructions, with the exception of using X065 assay buffer, and absorbance was measured at 450 nm on a BioTek 800 TS plate reader. Final concentrations (in ng/mL) were calculated from the standard curves on MyAssays.com and adjusted for the appropriate dilution factor before being compared among groups using Anova followed by Tukey post hoc contrasts.

#### iii. RNA extraction

RNA was extracted using Qiagen RNeasy® Plus Mini kits (Qiagen, Venlo, The Netherlands) following standard protocols for purification of RNA from animal tissues. Briefly, we added 350 μL of 0.5% Reagent DX-RLT lysis buffer to cryotubes containing frozen brains and completely homogenized tissue using a beadmill homogenizer (TissueLyser®, Qiagen, Venlo, The Netherlands) for two minutes at a time, flipping tubes once (four minutes total). We followed manufacturer instructions to elute RNA in 30 μL RNase-free water. Total RNA concentrations of extractions were quantified on a Qubit (Invitrogen, Waltham, MA, USA), and RNA was stored at

-80°C until further processing.

#### iv. Transcriptome Assembly, Annotation, and Transcript Quantification

Quality control, RNA library construction, and sequencing were performed at the Roy J. Carver Biotechnology Center at UIUC. Libraries were prepared with the Kapa Hyper Stranded mRNA library kit (Roche Diagnostics Corporation, Indianapolis, IN, USA). Sequencing was carried out on an Illumina Nova X Plus platform with V1.0 sequencing kits and a 150 nt paired-end protocol, yielding on average >65 M reads per sample. Transcriptome assembly and processing was performed on the TinkerCliffs computing cluster supported by the Advanced Research Computing unit at Virginia Tech University (Appendix D). Briefly, a *de novo* assembly was constructed and annotated from corrected paired-end reads following the Trinity-Trinotate pipeline [38,39]. On average 93.44% of reads mapped back to the final, filtered assembly. For downstream analyses, transcript abundances for each sample were aggregated into a single count matrix at the gene level using Trinity’s gene_counts_matrix, which yielded a total 859,656 “genes” of which 43,820 were annotated.

#### v. Differential Expression Analysis

Differential expression analysis was carried out in R Studio v.3.386. We compared gene expression across sexes and behavioral groups using DESeq2 [40]. For each group, low-count genes (<10 in at least half of samples) were filtered prior to analysis, yielding 65,000 to 75,000 genes per group. We filtered each group separately to maintain the maximum number of genes for each comparison. To visualize variation between groups, raw expression data was transformed using the variance stabilizing transformation and plotted using the plotPCA function. Principal component scores obtained from this analysis were compared between behavioral groups with Anova followed by Tukey post hoc comparisons. We estimated log-fold changes in expression using the ‘ashr’ shrinkage estimator [41] with a false discovery rate cutoff of α=0.05 applied to all P-values.

To investigate the function of transcriptomic differences across behavioral groups, we queried sets of genes differentially expressed at α=0.1 in each pairwise comparison for enriched GO terms. We used topGO to search for annotations in all three GO categories: biological processes, molecular function, and cellular component [42]. The nodeSize parameter was set to 10 to remove GO terms with fewer than 10 annotated genes and only terms with >1 significant gene were retained. Significance was estimated using a Fisher’s exact test.

## Results

### Male Call Rates Vary Across Parenting Cycle and Between Social Conditions

Overall, males called at significantly higher rates in trials where females were present (94.96 ± 42.51 calls/hour) compared to trials where females were removed (44.56 ± 37.13 calls/hour; 𝒳^2^ = 62.013, *P* < 0.0001; Fig. 1). Call rates also varied across stages of parenting (𝒳^2^ = 29.567, *P* < 0.0001): call rates were higher after eggs hatched but prior to tadpole transport in both treatments (Female Removal: Tukey HSD = 4.959, *P* < 0.0001; Control: Tukey HSD = 5.436, *P* < 0.0001), and in control trials (both partners present) call rates remained elevated during tadpole transport (Tukey HSD = 2.987, *P* = 0.0336). In female removal trials, call rates decreased between the pre-transport and transport stages (Tukey HSD = 6.336, *P* < 0.0001).

**Figure 1:**
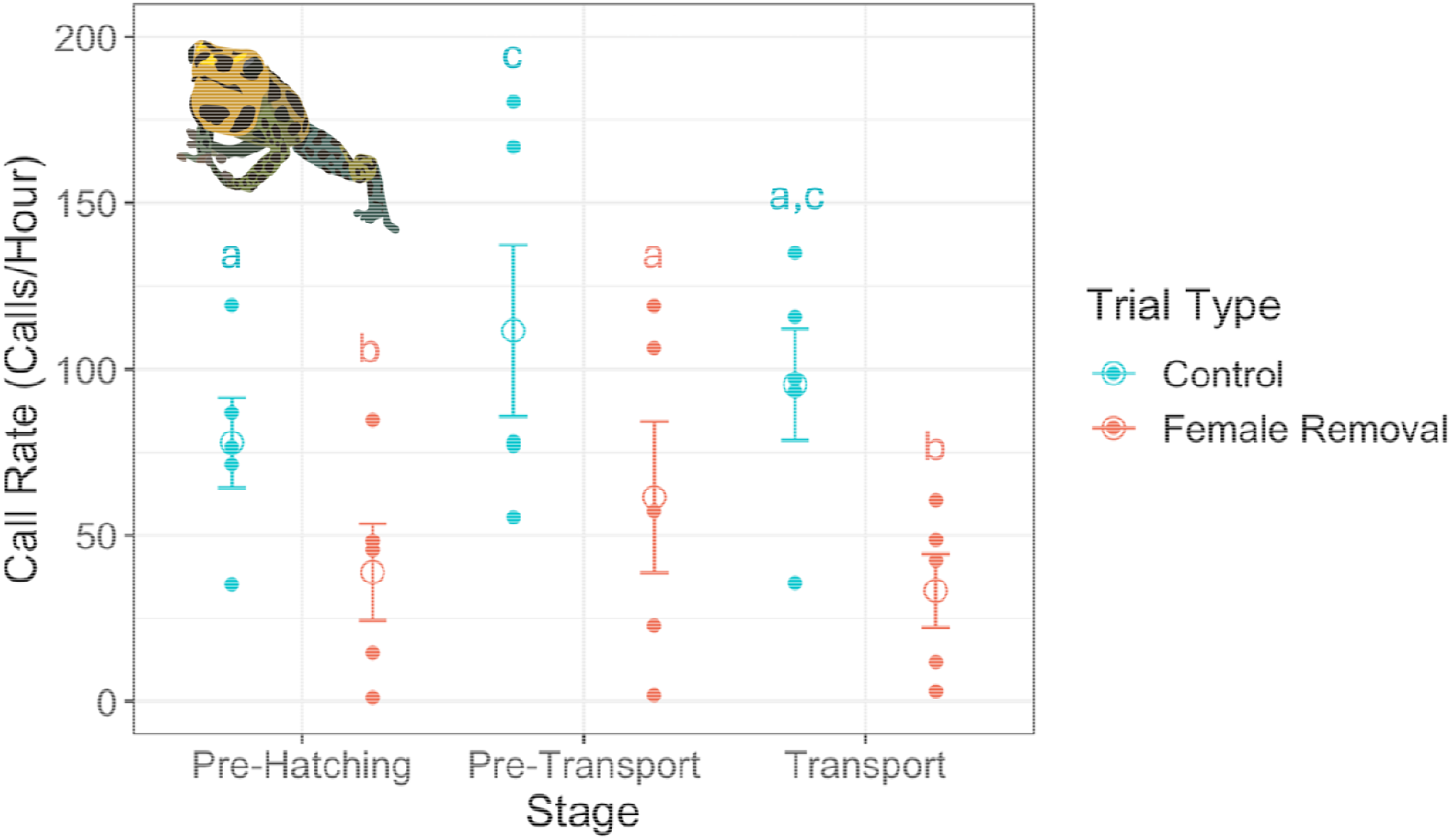
Variation in male call rates by trial type (female present, i.e., ‘Control’, or female removed). Asterisks denote significant departures from pre-hatching call rates observed in control trials (in blue) and female removed trials (in coral); Letters distinguish statistically significant differences between groups in pairwise comparisons.

### Sex and Treatment Differences in Transport Success

Transporting sex accounted for significant variation in transport success across trials (^2^ = 9.541, *P* = 0.002; Fig. 2). Males transported tadpoles in >95% of Control (i.e., both males and females were present) and Female Removal trials, but in trials where males were removed and transport could only be performed by females, roughly half of trials resulted in failure. We predicted that exposing females to acoustic and visual cues emulating their partner would lead to lower transport rates compared to trials where these cues were absent. Consistent with this, females took over transport in at least half of male removal (i.e., male visual and acoustic cues absent; 58%) and visual only trials (i.e., male acoustic cues absent; 60%) but success rates declined in the presence of acoustic playback (40%). When females were exposed to both playback and visual cues, transport occurred in only 20% of trials (Fig. 2). However, differences between trial types were not statistically significant after accounting for transporting sex and individual ID (^2^ = 4.380, *P* = 0.357). We found no significant effects of trial order on transport success, either independently (^2^ = 1.555, *P* = 0.212) or in interaction with transporting sex (^2^ = 0.0002, *P* = 0.989). In sex-reversed trials, a female’s initial success in taking over transport had no significant impact on subsequent successes (^2^ = 0.733, *P* = 0.392).

**Figure 2:**
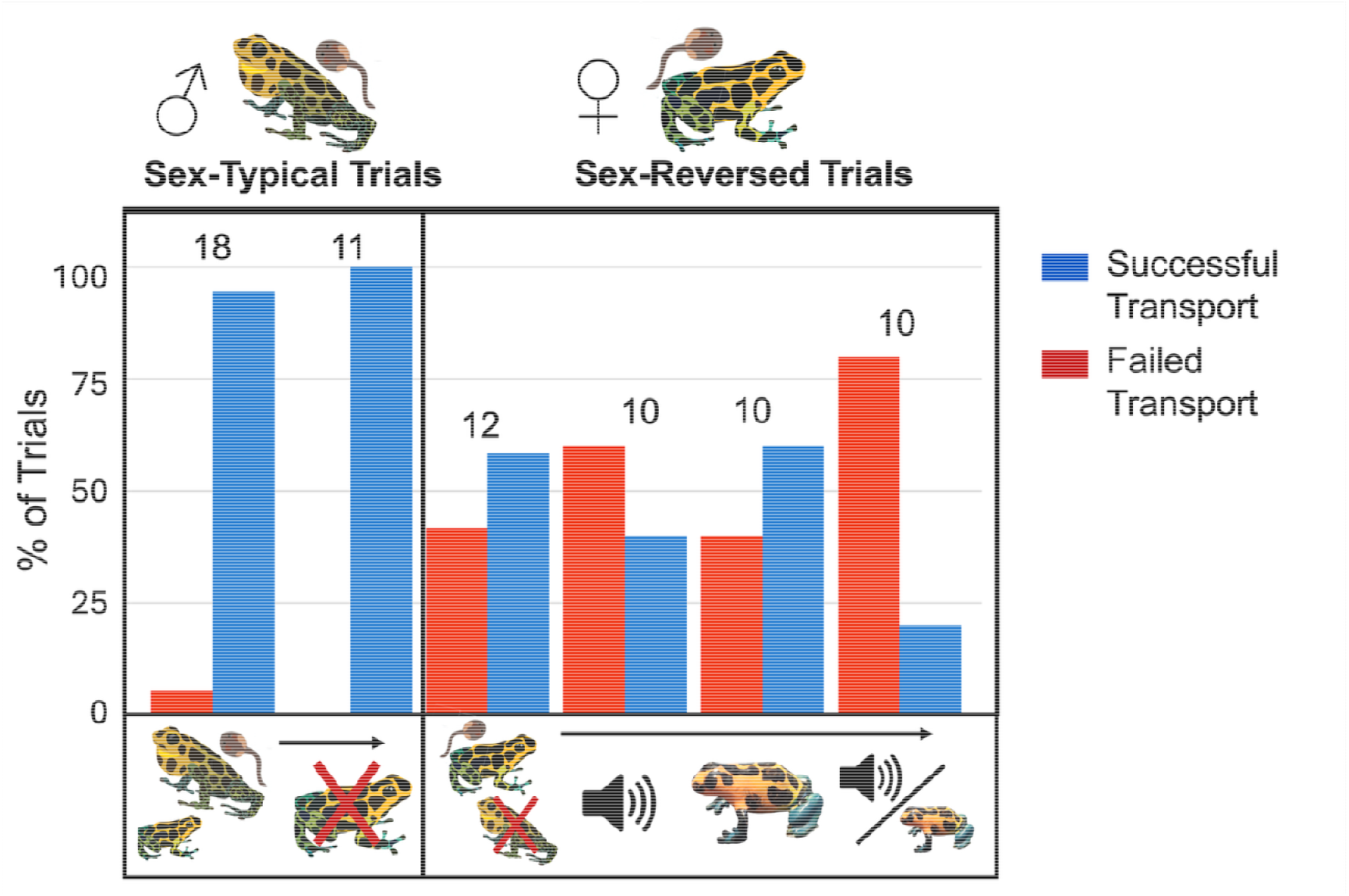
Variation in transport success across six trial types (from left to right): Control (both males and females present, no supplemental cues); Female removed (no supplemental cues); Male removed (no supplemental cues); Playback only (Male removed, advertisement calls broadcast into tank); Visual only (Male removed, dummy male situated near egg clutch); and Playback + Visual (Male removed, dumm male situated near egg clutch and advertisement cues broadcast into tank). Rates of transport success (blue) and failure (red) are depicted as a percentage of the total trials of that type, with number of trials denoted in text above the bars. In all sex-typical trials, transport was performed by the male, whether females were present or absent. In all sex-reversed trials, males were removed, and transport was performed by the female. In the one instance of transport failure observed in a sex-typical (Control) trial, the tadpole was picked up but fell off prior to deposition. In all other instances of transport failure, transport was not initiated.

Transport success was repeatable within individuals (R = 0.377, *P* = 0.015), a result that held when the analysis was restricted to only females and adjusted for trial type (R = 0.313, *P* = 0.061), meaning individuals who failed to transport in one trial type were more likely to also fail in a different trial type (Fig. 3).

**Figure 3:**
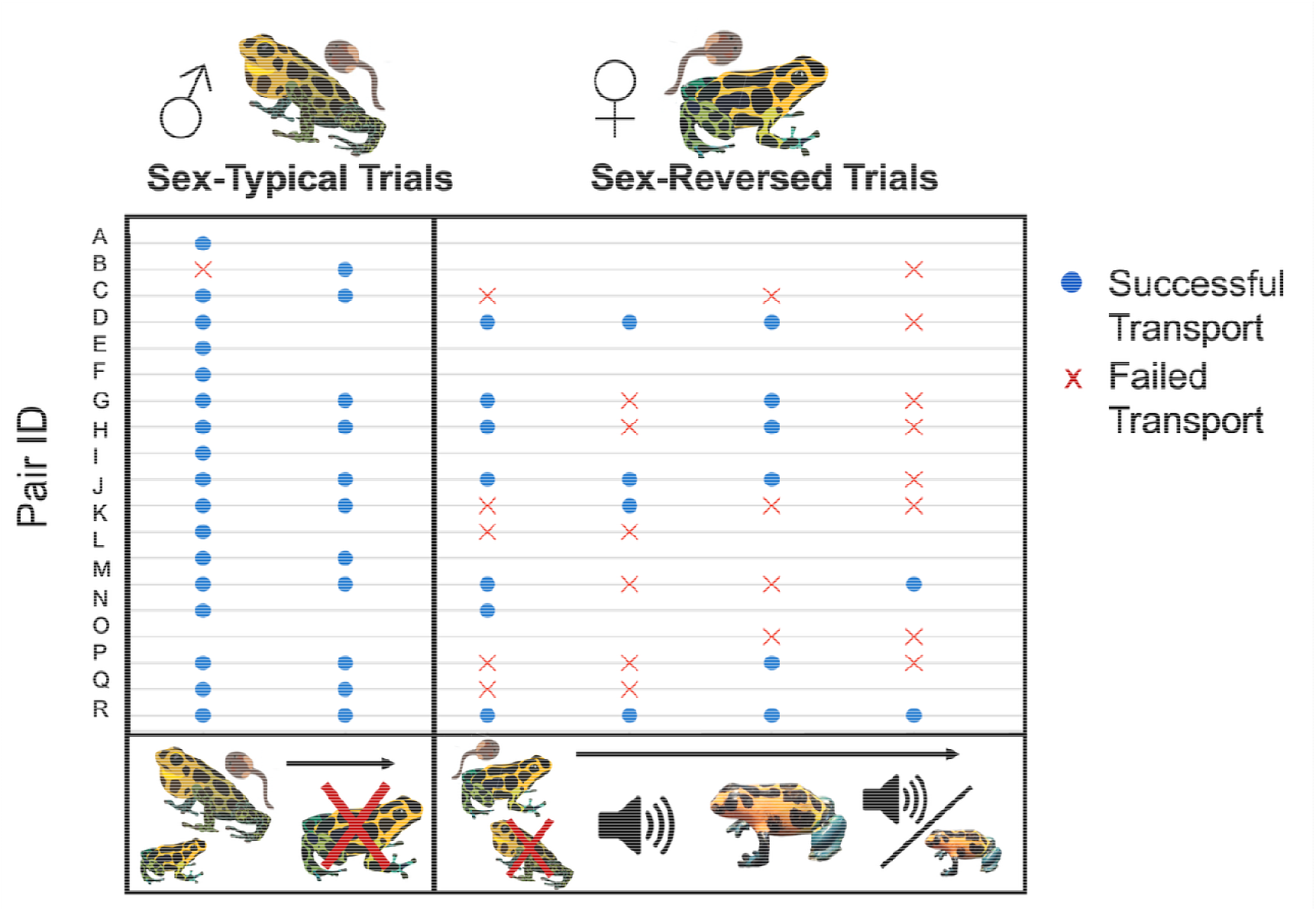
Individual variation in transport propensity across six trial types (from left to right): Control (both males and females present, no supplemental cues); Female removed (no supplemental cues); Male removed (no supplemental cues); Playback only (Male removed, advertisement calls broadcast into tank); Visual only (Male removed, dummy male situated near egg clutch); and Playback + Visual (Male removed, dummy male situated near egg clutch and advertisement cues broadcast into tank). Each row represents a distinct male-female pair, with male behavior summarized in the first two columns (sex-typical trials) and female behavior in the last four columns (sex-reversed trials). Blue circles denote successful transport, and red crosses denote failure of transport. Order of trials was randomized across pairs; however, not all individuals experienced all trial types.

### No Sex Differences in Transport Performance or Prehatching Behavior

Latency to transport did not differ significantly between males (0.826 ± 1.230 days; N=24) and females (1.467 ± 1.885 days; N=14) that transported tadpoles (Likelihood ratio test = 1.61, *P =* 0.211; Fig. S2A). Nor were there sex differences in transit time from tadpole pickup to drop-off (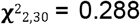, *P* = 0.592; Fig. S2B). Due to small numbers of transporting females in cue manipulation trials (i.e., as few as two), we do not report on differences in transport behavior across trial types.

All pre-hatching behaviors except for total time with eggs were significantly repeatable within individuals (number of pool visits: R=0.383, *P*=0.001; number of egg visits: R=0.178, *P*=0.050; total time in pool: R=0.251, *P*=0.018). No differences were observed between sexes or across treatments (*P* >0.05; Fig. S3–S4); however, some pre-hatching behaviors were predictive of post-hatching transport behaviors. Specifically, females that made more visits to eggs on the day before hatching were marginally more likely to transport tadpoles (𝒳^2^ = 2.811, *P* = 0.094; Fig. S5C). Among frogs that successfully transported tadpoles, males, but not females, that made more egg visitations on the day before hatching took less time to pick up hatched tadpoles (𝒳^2^ = 3.232, *P* = 0.072; Fig. S6A). In both sexes, shorter tadpole transit times were associated with higher frequency of prehatching pool visits (𝒳^2^ = 4.668, *P* = 0.031; Fig. S6B) and egg visits (𝒳^2^ = 15.943, *P* < 0.0001; Fig. S6C), with the latter effect more pronounced in female transporters (𝒳^2^ = 8.014, *P* = 0.005).

### Hormone Levels Sustained Across Parenting Cycle

Low plasma yields due to *R. imitator*’s small body size limited sample sizes for hormone comparisons. We found no significant differences in plasma cortisol across treatment groups (F_5,18_ = 1.417, *P* = 0.265) but note a non-significant trend of reduced cortisol levels in inflexible females compared to all other behavioral groups (Fig. 4; Tukey t = -1.73 – -2.58; *P* > 0.05).

**Figure 4:**
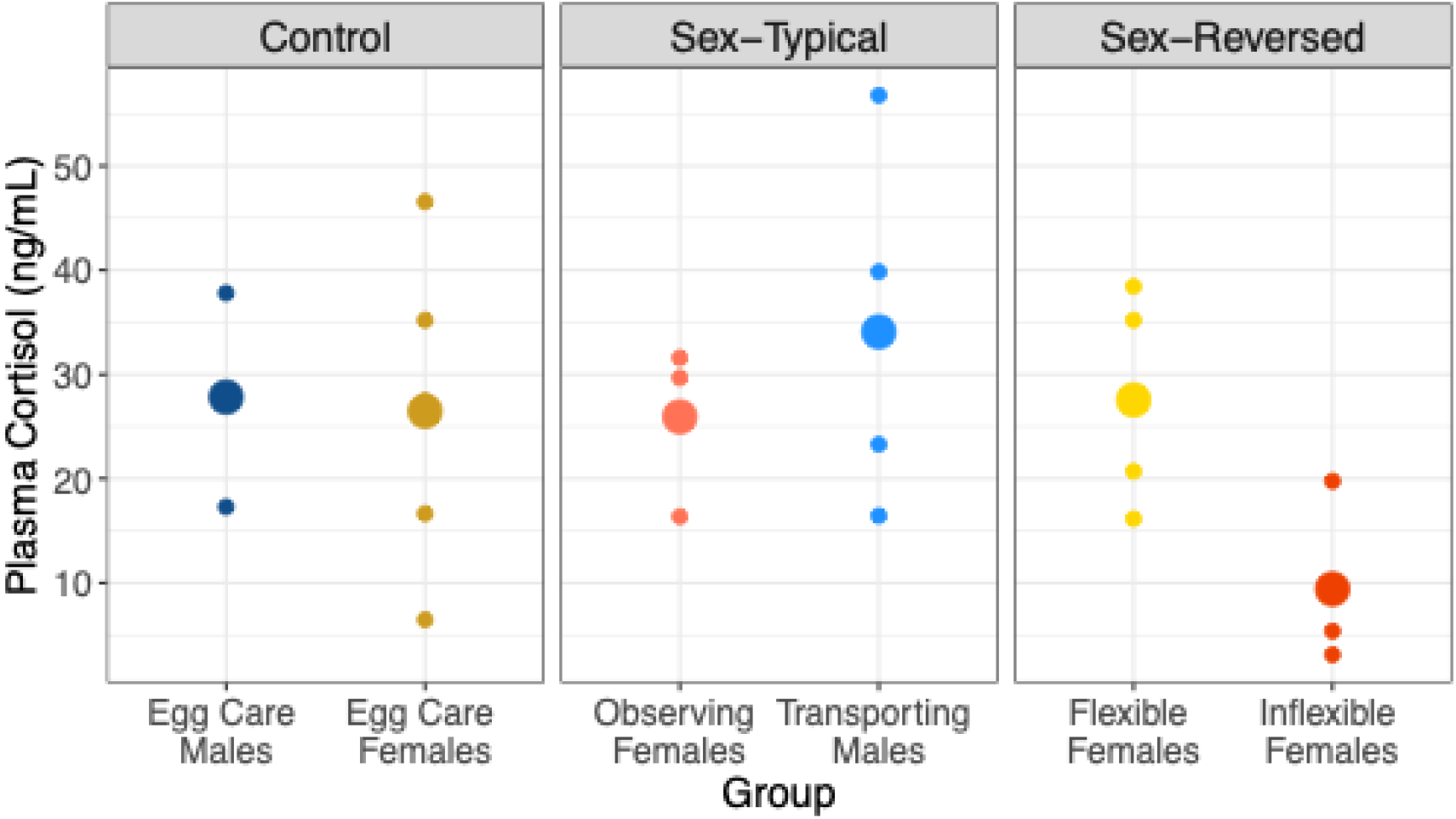
Concentrations of circulating cortisol, in ng/mL, across behavioral groups. Individuals were sacrificed at three timepoints: (1) during egg care, but before hatching (Egg Care Males and Egg Care Females); (2) after eggs hatched and tadpole transport was initiated (within one day of hatching; Observing Females, Transporting Males, and Flexible (transporting) Females); and (3) after tadpoles had been hatched for three days without transport (Inflexible (non-transporting) Females). Small points denote individual samples and large points denote group means.

### Individual Variation in Plasticity Associated with Gene Expression Signature

We examined differences in brain gene expression associated with differences in parental behavior using PCA. Samples separated along two primary principal components, which cumulatively explained 29% (PC1=17.9%, PC2=11.1%) of variance in overall gene expression (Fig. 5). Differences among groups were marginally significant when considering separation along PC1 (F_5,24_=2.116; *P*=0.098), but not PC2 (F_5,24_=0.6915; *P*=0.635). In sex-typical trials, egg care males, egg care females, transporting males, and observing females exhibited overlapping gene expression profiles, with no significant separation among groups (PC1: F_5,24_=0.437; *P*=0.730; PC2: F_5,24_=0.953; *P*=0.440; Fig. 5A). In sex-reversed trials, flexible (transporting) females overlapped with egg care females, whereas inflexible (non-transporting) females separated significantly from caregiving parents along PC1 (F_5,24_=6.401; *P*=0.013; Egg Care– Non-transporting: Tukey=19.933, *P*=0.029; Transporting–Non-transporting: Tukey=19.292, *P*=0.020; Fig. 5B).

**Figure 5:**
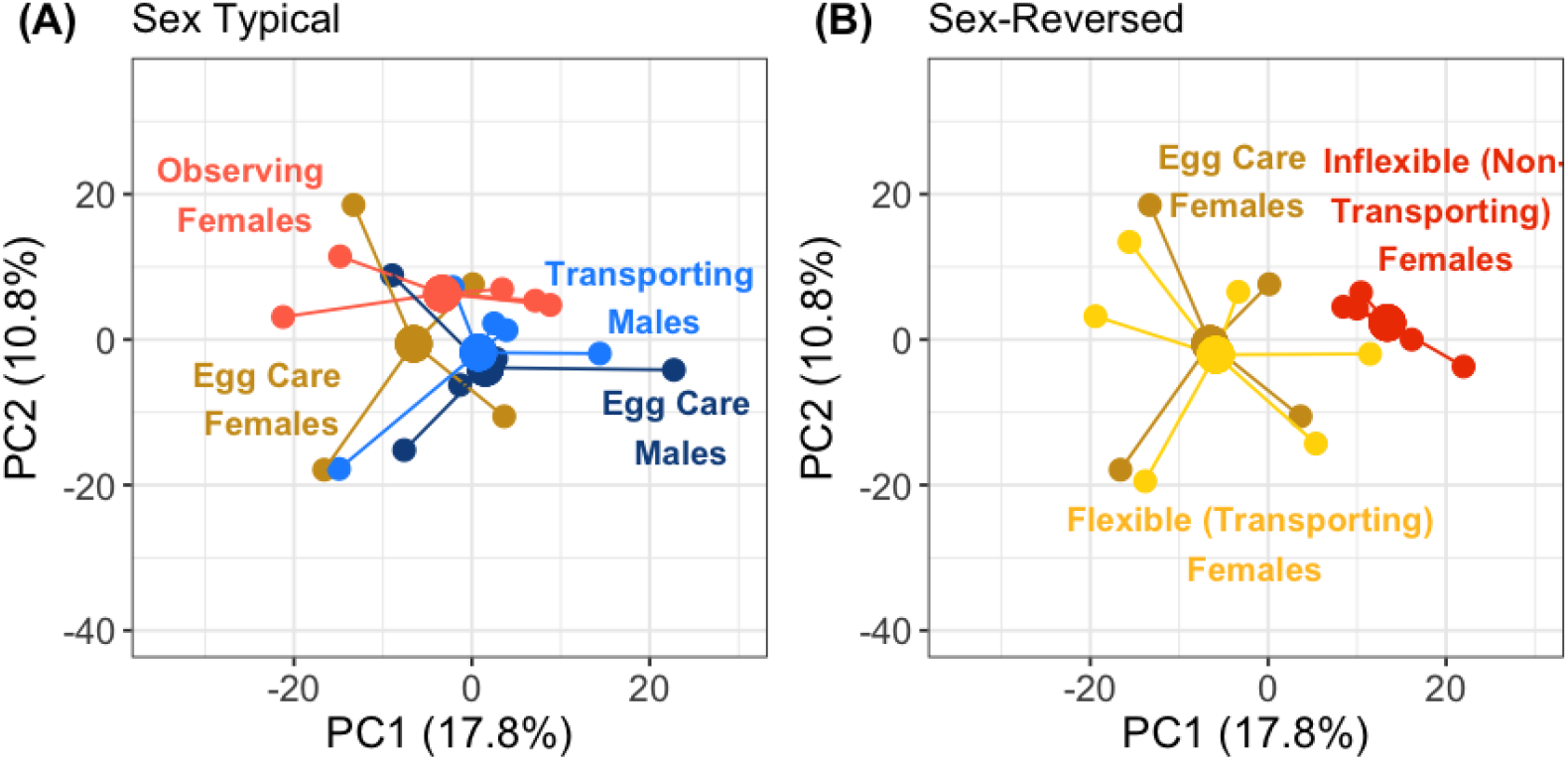
Principal components analysis of brain gene expression patterns among behavioral groups belonging to sex-typical trials (on the left) and sex-reversed trials (on the right). Same-sex egg care samples are plotted in both figures for reference.

The overall distribution of gene expression profiles was underscored by patterns of differential gene expression between groups. Relative to egg care parents of the same sex, transporting males exhibited more differences in gene expression (N=74 differentially expressed genes; Table S2) than either observing (N=54 differentially expressed genes; Table S3) or transporting females (N=23 differentially expressed genes; Fig. S7; Table S4). Transporting males and females differed significantly in the expression of N=14 genes (Table S5). The highest number of differentially expressed genes of any group was associated with females that failed to transport, which arose when comparing to both egg care (N=178; Fig. S7; Table S6) and transporting females (N=164; Table S7). There were 69 genes overlapping between these two contrasts (Tables S5–6), among them enzymes involved in the hydrolysis of peptide and steroid hormones (e.g., Mephrin A subunit alpha, aldo-keto reductase family 1 member C4) and the synthesis and inhibition of neurotransmitters (e.g., Gamma-aminobutyric acid receptor subunit rho-1, Pterin-4-alpha-carbinolamine dehydratase). Butyrophilin subfamily 3 member A3 and E3 ubiquitin/ISG15 ligase, two immune genes that were significantly downregulated in transporting females relative to egg care females (Butyrophilin subfamily 3 member A3: log2fold change=-7.957; E3 ubiquitin/ISG15 ligase: log2fold change=-7.959; Table S4), were also downregulated in the contrast between transporting and non-transporting females (Butyrophilin subfamily 3 member A3: log2fold change=-8.417; E3 ubiquitin/ISG15 ligase: log2fold change=-6.904; Table S7).

Within the transport stage, a female’s social state (i.e., presence or absence of male partner) also accounted for differences in gene expression (N=94 differentially expressed genes), notably in genes related to circulation (e.g., calcitonin gene-related peptide type 1 receptor isoform X1, histidine-rich glycoprotein-like) and neuronal function (e.g., Glycogen phosphorylase (brain form), synaptotagmin-17 isoform X3, tectonin beta-propeller repeat-containing protein 2, pumilio homolog 2 isoform X2; Table S8).

Inspection of GO annotations across differentially expressed genes revealed significant enrichment of immune response categories in transporting males (e.g., GO:0048525, negative regulation of viral process, GO:0050792, regulation of viral process, GO:0032897, negative regulation of viral transcription; Table S9) and transporting females (e.g., GO:0002757, immune response-activating signaling pathway, GO:0002764, immune response-regulating signaling pathway, GO:0002253, activation of immune response; Table S10). Similarly, the major pathways enriched in females observing male transport involved proliferation of T cells (e.g., GO:0046006, regulation of activated T cell proliferation, GO:0050798, activated T cell proliferation, GO:0043382, positive regulation of memory T cell differentiation) and other immune cells (GO:0045621, positive regulation of lymphocyte differentiation, GO:0070663, regulation of leukocyte proliferation, GO:0032604, granulocyte macrophage colony-stimulating factor production; Table S11–S12).

Finally, genes differentially expressed between inflexible, non-transporting females and flexible, transporting females were significantly enriched in functions related to tissue repair (e.g., GO:0060155, platelet dense granule organization, GO:0048041, focal adhesion assembly, GO:0007044, cell-substrate junction assembly) and homeostasis (e.g., GO:2001137, positive regulation of endocytic recycling; GO:2001135, regulation of endocytic recycling; Table S14).

## Discussion

Flexible parenting is widespread in nature and often differs between the sexes. Manipulations aimed at identifying the mechanisms responsible for the induction of care in the typically non-caregiving sex can offer critical insights into the processes that promote cross-sexual transfer and the evolutionary diversification of parental care between species [1,3]. Here, we studied the extrinsic and intrinsic factors affecting sex-reversed tadpole transport in the biparental poison frog, *Ranitomeya imitator*. By manipulating social conditions experienced by parents of both sexes, we exposed sex and individual variation in the propensity to perform a typically male-biased parental behavior. Collectively, our findings suggest that sex-reversed (female) tadpole transport is influenced by male signaling, that individual females vary in the propensity for tadpole transport, and that maintaining patterns of gene expression and hormones associated with a caring state may be a prerequisite to the takeover of transport behavior.

### (a) The sexes differ predictably in parental behavior and propensity to perform a specific task

The first goal of this study was to characterize sex and individual variation in tadpole transport behavior in a biparental frog for which sex-reversed care (i.e., female tadpole transport) has been documented but not systematically studied. We predicted that if the necessity for female tadpole transport arises infrequently in nature, then both the propensity and performance of this task should be male-biased, whereas shared tasks should not show such sex bias. Consistent with previous behavioral accounts of *R. imitator* [25], we found that males and females contribute equally to pre-hatching egg care. However, we observed significant sex differences in transport success, with males successfully transporting tadpoles in 96.6% of trials and females transporting in only 50% of trials when males were removed, supporting our prediction that females would underperform in a typically male-biased task. Females in our trials were never observed to takeover transport in the presence of males, although we have occasionally observed this behavior in the lab other contexts.

The confirmation that a strong sex bias in tadpole transport behavior prevails while both parents are present implies that either males consistently pick up and transport tadpoles before females can or that females actively defer to their male partners. The former explanation seems unlikely, as we observed no sex differences in the latency to pick up tadpoles. In addition, *R. imitator* typically lay 2–4 eggs but transport only one tadpole at a time, such that unavailability of hatched tadpoles is unlikely to limit female transport in the majority of circumstances. We also could not attribute transport failures to tadpoles being lost in transit, as this was observed in only a single instance where the transporter was a male (Fig. 2). Rather, variation in transport success across trials was explained by variation in transport *propensity*, with females less likely than males to initiate transport in all social conditions. Post-pickup, male and female transport behavior (i.e., transit time from pickup to drop-off) was indistinguishable, suggesting that intrinsic sex differences affect the probability of females initiating transport, but not transport performance. Intriguingly, individual variation in female behavior prior to egg hatching predicted transport propensity in male removal trials. Specifically, females that took over transport made more frequent visitations to eggs on the day before hatching. Although this effect was only marginally significant, it supports the idea that female decisions about transport may precede the window for transport. Thus, environmental and social conditions experienced in earlier stages of care should influence these decisions.

### (b) Partner cues influence probability of takeover by the non-typical caregiver

Results of our mate removal and cue manipulation experiments offer further insights into the extrinsic factors shaping sex-specific parental behavior. Our study provides two lines of evidence supporting the prediction of Ringler *et al*., (2015), that females monitor male acoustic signals to inform takeover decisions. We previously showed that pair-bonded female *R. imitator* discriminate and preferentially associate with advertisement calls of their partner [31], and here we show that male *R. imitator* modify calling behavior in the absence of female partners and across care stages. While continuous calling activity from within one’s territory likely function in the maintenance of territorial boundaries during care (i.e., male-male communication; [23,29,44]), results of our female removal trials imply an additional role in within-pair communication.

Clues as to the precise function for this calling behavior come from our male removal trials, in which females with mates removed were provided audio recordings of their partner’s calls. Of the six repeat-assayed females that successfully took over transport in standard mate removal experiments, three failed to transport in the presence of audio playback alone and four did not transport when presented with both playback and a visual cue. While this result did not attain statistical significance due to sample size, these patterns are consistent with the expectation that females are attuned to partner cues and use these to make care decisions [28]. Similar patterns have been demonstrated in other biparental species, whereby one sex disproportionately ‘monitors’ efforts by the other to inform compensatory responses (e.g., [45]). While such social cues may be broadcast inadvertently and used in place of offspring cues to minimize effort in direct evaluation [46,47], in the case of *R. imitator* it appears that males intentionally broadcast acoustic signals to females and even modulate calling effort in anticipation of transport, consistent with other lines of evidence that implicate acoustic communication in the coordination biparental care [27]. The absence of any sex or treatment differences in egg visitation rates in our study further disqualifies the possibility that females use male cues as a substitute for offspring cues. Rather, it seems that females adhere to a simple decision rule of refraining from tadpole transport if males are present, and that the acoustic and visual cues deployed in our experiments serve as placeholders (albeit imperfect ones) for an absent partner.

### (c) Individual variation in transport propensity is repeatable

Another key result of our behavioral experiment was the demonstration of repeatable individual variation in parenting behaviors. Indeed, both prehatching behaviors and transport success were significantly repeatable within individuals, and females that failed to transport in the standard mate removal trial also failed to transport in a majority of cue manipulation trials (Fig. 3). The extent of individual variation in parenting behavior in poison frogs and the factors that account for this variation are generally poorly understood. In our study, the propensity for sex-reversed tadpole transport did not reflect age, parental experience, or morph, as subjects classed as ‘inflexible’ spanned the full range of these factors (Table S1). Regardless of causal factors, individual variation in care can have important evolutionary implications. A recent study involving individual-based evolutionary simulations showed that transient, within-sex polymorphisms in care behavior are almost always the first step in the evolution of sex role specialization [8]. Indeed, individual variation is theorized to be a critical substrate in the evolutionary diversification of parenting systems *via* cross-sexual transfer of parental behaviors [3]. While the conditions that expose polymorphisms in female transport ability are expected to arise relatively infrequently in *R. imitator* due to the high degree of male coordination that is necessary across stages of care [24,27], the existence of repeatable individual variation nonetheless entertains the possibility that a sudden shift in conditions could promote directional selection for females that provide care.

### (d) The transition to active parenting is associated with subtle changes in gene expression

Consistent with previous hormonal and neurogenomic investigations of parental care in dendrobatids (e.g., [12,32]), we observed minimal sex differentiation in circulating cortisol or brain gene expression. Remarkably, when comparing parents in the egg care stage to parents in the transport stage, neither sex showed large changes in cortisol concentrations or gene expression whether in the sex-typical condition (male transporting, female observing) or sex-reversed condition (female transporting, male absent). Our results differ from those of Fischer & O’Connell (2020), who in *D. tinctorius* observed significant increases in circulating cortisol and significantly altered expression of hundreds to thousands of transcripts in both sexes in association with the transition to active transport. Species differences may account for some of this variation; for example, because *R. imitator* is a pair bonding species, changes in hormone circulation and gene expression may arise early in the cycle to synchronize the physiological and neural states of males and females (e.g., [48]). Indeed, large changes in neural induction in *R. imitator* when comparing non-parental frogs to parents in the transport stage [32] support the existence of an overall ‘caring state,’ of which many aspects may persist across care stages.

While subtle, variation in brain gene expression detected with our sampling design nevertheless revealed familiar mechanisms associated with parental care, including enrichment of immune response pathways in both observing and actively transporting parents relative to egg care parents. Interestingly, greater distinctions in neurogenomic state arose when comparing females in the presence (observing) *versus* absence (transporting and non-transporting) of male partners within a care stage than between care stages or behavioral states, most notably in genes related to circulation and neuronal function. This implies that mate removal triggers changes in female neural states regardless of subsequent behavioral decisions, an effect which may be mediated in part by male acoustic signaling.

### (e) Absence of behavioral plasticity is associated with shifts in neurogenomic state

Contrary to our prediction that large changes in gene expression would be associated with the activation of tadpole transport behavior, the most profound differences in our study consistently arose from contrasts involving females identified *a priori* as inflexible, on the basis of having previously failed to takeover transport. While our behavioral analyses showed that transport propensity is repeatable, to ensure that females were not sampled prematurely (i.e., before having the opportunity to transport) sampling was postponed a day beyond the 80% transport window. Inflexible females showed non-significant reductions in circulating cortisol and significantly altered expression of genes involved in hormone and neurotransmitter regulation compared to egg caring females. Additional differences included the upregulation of immune genes that were significantly downregulated in flexible females during transport. Given that many expression differences attributed to non-transporting females were shared between contrasts with egg care and transporting females, we suggest that the neurogenomic state occupied by inflexible females represents a departure not only from a transport-capable state, but from a caring state altogether. There are at least two possible interpretations of these patterns: (1) inflexible females never achieve a caring state to begin with, and this is the cause of their failure to transport; (2) inflexible females prematurely exit the caring state, potentially in response to an environmental trigger such as mate removal. Distinguishing between these alternatives will require further experimentation using designs with non-parental adult controls and sampling across multiple time points. However, if male removal does serve as the trigger for inflexible females ‘exiting’ a caring state, it would suggest that the same environmental stimulus induces opposite responses in different females, consistent with the existence of a cryptic within-sex polymorphism.

## Conclusion

This study identified sex differences in the extrinsic motivators, but not the underlying neurogenomic mechanisms, of a sex-biased parental behavior in a poison frog with clear division of labor, providing insights into how simple decision rules reinforced by communication within pairs can both maintain typical sex roles and promote flexibility. Specifically, we show that male calling behavior can be used to track the progression of parenting cycles from egg care to tadpole transport and that these signals may function in suppressing parenting behaviors in females while males are present. Our data further suggest that the activation of specific parenting behaviors (here, tadpole transport) need not involve large changes in neurogenomic state; rather, genes expressed during ‘priming’ may account for the majority of what is needed for the active performance of the behavior, whether by males or females. This finding reinforces our behavioral data, which show that male and female transporters do not differ in any observable performance measure (e.g., latency to initiate transport or duration of transport) but rather differ only in propensity to transport. Importantly, individuals lacking the capacity for sex-reversed behavior show subtle variation in parental behaviors leading up to the window for transport and exhibit a distinct neurogenomic state during the transport window. Therefore, rather than being associated with dramatic changes in neurogenomic state in response to a social trigger, the mechanisms responsible for sex-reversed parental behavior appear to lie in the ability to access and maintain activation of shared machinery even under sex-typical conditions.

## Acknowledgements

We thank Mark Bee and James Tumulty for helpful advice on acoustic playback methodologies during early project design stages and the Champaign-Urbana Community Fab Lab for assistance with 3D printing. We thank the Fischer lab for helpful feedback.

## Funding

This study was supported by a National Science Foundation Postdoctoral Research Fellowship in Biology to J.B.M. (2010649), a National Science Foundation grant to E.K.F. (IOS 21-46058), a University of Illinois Urbana-Champaign Research Board Grant to E.K.F. (RB21025), University of Illinois Urbana-Champaign laboratory start-up funds to E.K.F., and Virginia Tech laboratory start-up funds to J.B.M.

## Notes

### Competing Interest Statement

The authors have declared no competing interest.

## References

1. West-Eberhard MJ. 2003 Developmental plasticity and evolution. Oxford, U.K.: Oxford University Press.

2. Anderson AP, Falk JJ. 2023 Cross-sexual Transfer Revisited. Integrative and Comparative Biology 63, 936–945. (doi:10.1093/icb/icad021)

3. Westrick SE, Moss JB, Fischer EK. 2023 Who cares? An integrative approach to understanding the evolution of behavioural plasticity in parental care. Animal Behaviour 200, 225–236.

4. Kokko H, Jennions MD. 2013 Sex differences in parental care. In The Evolution of Parental Care (eds NJ Royle, PT Smiseth), pp. 101–116. Oxford, UK: Oxford University Press. (doi:10.1093/acprof:oso/9780199692576.003.0006)

5. McNamara JM, Wolf M. 2015 Sexual conflict over parental care promotes the evolution of sex differences in care and the ability to care. Proceedings of the Royal Society B: Biological Sciences 282, 20142752. (doi:10.1098/rspb.2014.2752)

6. Fromhage L, Jennions MD. 2016 Coevolution of parental investment and sexually selected traits drives sex-role divergence. Nature Communications 7, 1–11. (doi:10.1038/ncomms12517)

7. Henshaw JM, Fromhage L, Jones AG. 2019 Sex roles and the evolution of parental care specialization. Proceedings of the Royal Society B: Biological Sciences (doi:10.1098/rspb.2019.1312)

8. Long X, Weissing FJ. 2023 Transient polymorphisms in parental care strategies drive divergence of sex roles. Nat Commun 14, 6805. (doi:10.1038/s41467-023-42607-6)

9. Royle NJ, Russell AF, Wilson AJ. 2014 The evolution of flexible parenting. Science 345, 776–781. (doi:10.1126/science.1253294)

10. Parker DJ, Cunningham CB, Walling CA, Stamper CE, Head ML, Roy-Zokan EM, McKinney EC, Ritchie MG, Moore AJ. 2015 Transcriptomes of parents identify parenting strategies and sexual conflict in a subsocial beetle. Nature Communications 6, 8449. (doi:10.1038/ncomms9449)

11. Kohl J et al. 2018 Functional circuit architecture underlying parental behaviour. Nature 556, 326–331. (doi:10.1038/s41586-018-0027-0)

12. Fischer EK, O’Connell LA. 2020 Hormonal and neural correlates of care in active versus observing poison frog parents. Hormones and Behavior 120, 104696. (doi:10.1016/j.yhbeh.2020.104696)

13. Lopes PC, de Bruijn R. 2021 Neurotranscriptomic changes associated with chick-directed parental care in adult non-reproductive Japanese quail. Scientific Reports 2021 11:1 11, 1– 11. (doi:10.1038/s41598-021-94927-6)

14. Roland AB, O’Connell LA. 2015 Poison frogs as a model system for studying the neurobiology of parental care. Current Opinion in Behavioral Sciences 6, 76–81. (doi:10.1016/j.cobeha.2015.10.002)

15. Furness AI, Capellini I. 2019 The evolution of parental care diversity in amphibians. Nature Communications 10, 4709.

16. Liedtke HC, Wiens JJ, Gomez-Mestre I. 2022 The evolution of reproductive modes and life cycles in amphibians. Nat Commun 13, 7039. (doi:10.1038/s41467-022-34474-4)

17. Ringler E, Rojas B, Stynoski JL, Schulte LM. 2023 What Amphibians Can Teach Us About the Evolution of Parental Care. Annual Review of Ecology, Evolution, and Systematics 54, 43–62. (doi:10.1146/annurev-ecolsys-102221-050519)

18. O’Connell LA. 2020 Frank Beach Award Winner: Lessons from poison frogs on ecological drivers of behavioral diversification. Hormones and Behavior 126, 104869. (doi:10.1016/j.yhbeh.2020.104869)

19. Summers K, Weigt LA, Boag P, Bermingham E. 1999 The evolution of female parental care in poison frogs of the genus Dendrobates: Evidence from mitochondrial DNA sequences. Herpetologica

20. Summers K, Tumulty J. 2014 Parental care, sexual selection, and mating systems in Neotropical poison frogs. In Sexual Selection: Perspectives and Models from the Neotropics, pp. 289–320. Academic Press.

21. Caldwell JP, Oliveira VRL de. 1999 Determinants of Biparental Care in the Spotted Poison Frog, Dendrobates vanzolinii (Anura: Dendrobatidae). Copeia 1999, 565–575. (doi:10.2307/1447590)

22. Brown JL, Twomey E, Morales V, Summers K. 2008 Phytotelm size in relation to parental care and mating strategies in two species of Peruvian poison frogs. Behaviour 145, 1139– 1165. (doi:10.1163/156853908785387647)

23. Brown JL, Morales V, Summers K. 2010 A key ecological trait drove the evolution of biparental care and monogamy in an amphibian. American Naturalist 175, 436–446. (doi:10.1086/650727)

24. Tumulty J, Morales V, Summers K. 2014 The biparental care hypothesis for the evolution of monogamy: Experimental evidence in an amphibian. Behavioral Ecology 25, 262–270. (doi:10.1093/beheco/art116)

25. Schulte LM, Summers K. 2021 Who cares for the eggs? Analysis of egg attendance behaviour in Ranitomeya imitator, a poison frog with biparental care. Behav. 159, 603–614. (doi:10.1163/1568539X-bja10142)

26. Schulte LM. 2014 Feeding or avoiding? Facultative egg feeding in a Peruvian poison frog (Ranitomeya variabilis). Ethology Ecology & Evolution 26, 58–68. (doi:10.1080/03949370.2013.850453)

27. Moss JB, Tumulty JP, Fischer EK. 2023 Evolution of acoustic signals associated with cooperative parental behavior in a poison frog. Proceedings of the National Academy of Sciences 120, e2218956120. (doi:10.1073/pnas.2218956120)

28. Ringler E, Pašukonis A, Fitch WT, Huber L, Hödl W, Ringler M. 2015 Flexible compensation of uniparental care: Female poison frogs take over when males disappear. Behavioral Ecology 26, 1219–1225. (doi:10.1093/beheco/arv069)

29. Mayer M, Schulte LM, Twomey E, Lötters S. 2014 Do male poison frogs respond to modified calls of a Müllerian mimic? Animal Behaviour 89, 45–51. (doi:10.1016/j.anbehav.2013.12.013)

30. Twomey E, Mayer M, Summers K. 2015 Intraspecific call variation in the mimic poison frog Ranitomeya imitator. Herpetologica 71, 252–259. (doi:10.1655/HERPETOLOGICA-D-15-00004)

31. Podraza ME, Moss JB, Fischer EK. 2024 Evidence for individual vocal recognition in a pair-bonding poison frog, Ranitomeya imitator. Journal of Experimental Biology 227, jeb246753. (doi:10.1242/jeb.246753)

32. Fischer EK, Roland AB, Moskowitz NA, Tapia EE, Summers K, Coloma LA, O’Connell LA. 2019 The neural basis of tadpole transport in poison frogs. Proceedings of the Royal Society B: Biological Sciences 286. (doi:10.1098/rspb.2019.1084)

33. Bates DM, Maechler M, Bolker B, Walker S. 2015 Fitting linear mixed-effects models using lme4. Journal of statistical software 67, 1–48. (doi:10.1088/1742-6596/43/1/292)

34. Lenth RV, Buerkner P, Herve M, Love J, Miguez F, Riebl H, Singmann H. 2022 Package ‘emmeans’: Estimated marginal means, aka least-squares means. R Package version 3.4.0 (doi:10.1080/00031305.1980.10483031)

35. Stoffel MA, Nakagawa S, Schielzeth H. 2017 rptR: repeatability estimation and variance decomposition by generalized linear mixed-effects models. Methods in Ecology and Evolution 8, 1639–1644. (doi:10.1111/2041-210X.12797)

36. Therneau TM, T. Lumley. 2015 Package ‘ survival ‘. R package version 3.2.7

37. Westrick SE, Paitz RT, Fischer EK. 2023 Why not both? A case study measuring cortisol and corticosterone in poison frogs., 2023.06.19.545597. (doi:10.1101/2023.06.19.545597)

38. Grabherr MG et al. 2011 Trinity: reconstructing a full-length transcriptome without a genome from RNA-Seq data. Nat Biotechnol 29, 644–652. (doi:10.1038/nbt.1883)

39. Bryant DM et al. 2017 A tissue-mapped axolotl de novo transcriptome enables identification of limb regeneration factors. Cell Rep 18, 762–776. (doi:10.1016/j.celrep.2016.12.063)

40. Love MI, Huber W, Anders S. 2014 Moderated estimation of fold change and dispersion for RNA-seq data with DESeq2. Genome Biology 15, 550. (doi:10.1186/s13059-014-0550-8)

41. Stephens M. 2017 False discovery rates: A new deal. Biostatistics 18, 275–294. (doi:10.1093/biostatistics/kxw041)

42. Alexa A, Rahnenfuhrer J. 2020 topGO: Enrichment analysis for gene ontology. R package version 2.42.0

43. Bukhari SA, Saul MC, James N, Bensky MK, Stein LR, Trapp R, Bell AM. 2019 Neurogenomic insights into paternal care and its relation to territorial aggression. Nature Communications 2019 10:1 10, 1–11. (doi:10.1038/s41467-019-12212-7)

44. Brown JL, Morales V, Summers K. 2009 Home range size and location in relation to reproductive resources in poison frogs (Dendrobatidae): a Monte Carlo approach using GIS data. Animal Behaviour 77, 547–554. (doi:10.1016/j.anbehav.2008.10.002)

45. Cotter SC, Kilner RM. 2010 Sexual division of antibacterial resource defence in breeding burying beetles, Nicrophorus vespilloides. Journal of Animal Ecology 79, 35–43. (doi:10.1111/j.1365-2656.2009.01593.x)

46. Danchin É, Giraldeau LA, Valone TJ, Wagner RH. 2004 Public information: From nosy neighbors to cultural evolution. Science 305, 487–491. (doi:10.1126/science.1098254)

47. Ringler E, Pašukonis A, Ringler M, Huber L. 2016 Sex-specific offspring discrimination reflects respective risks and costs of misdirected care in a poison frog. Animal Behaviour 114, 173–179. (doi:10.1016/j.anbehav.2016.02.008)

48. Nowicki JP, Rodríguez C, Lee JC, Goolsby BC, Yang C, Cleland TA, O’Connell LA. 2024 Physiological state matching in a pair bonded poison frog. Royal Society Open Science 11, 240744. (doi:10.1098/rsos.240744)

