## Appendix A, Appendix B, Appendix C, Appendix D for "Partner cues and individual variation underlie sex-reversed parental care in poison frogs"

*Supplementary Methods*

A. *Visual Stimuli*

To create realistic dummy frogs to serve as visual stimuli for this experiment, plastic models were printed to scale (17x12x9 mm LxWxH) by modifying a published .STL file (https://www.printables.com/en/model/211953-poison-dart-frog) in Tinkercad (Autodesk, Inc., San Francisco, CA). 3D-printing was performed on a Flashforge Finder FDM 3D printer (Zhejiang Flashforge 3D technology Co., LTD, Zhejiang, China) at the Champaign-Urbana Community Fab Lab. To maximize resemblance to focal males, models were painted from reference photos to match individual color patterns using modeling paints (Fig. 2). Models were finished with glosscote spray to give a reflective appearance and mounted on suction cups.

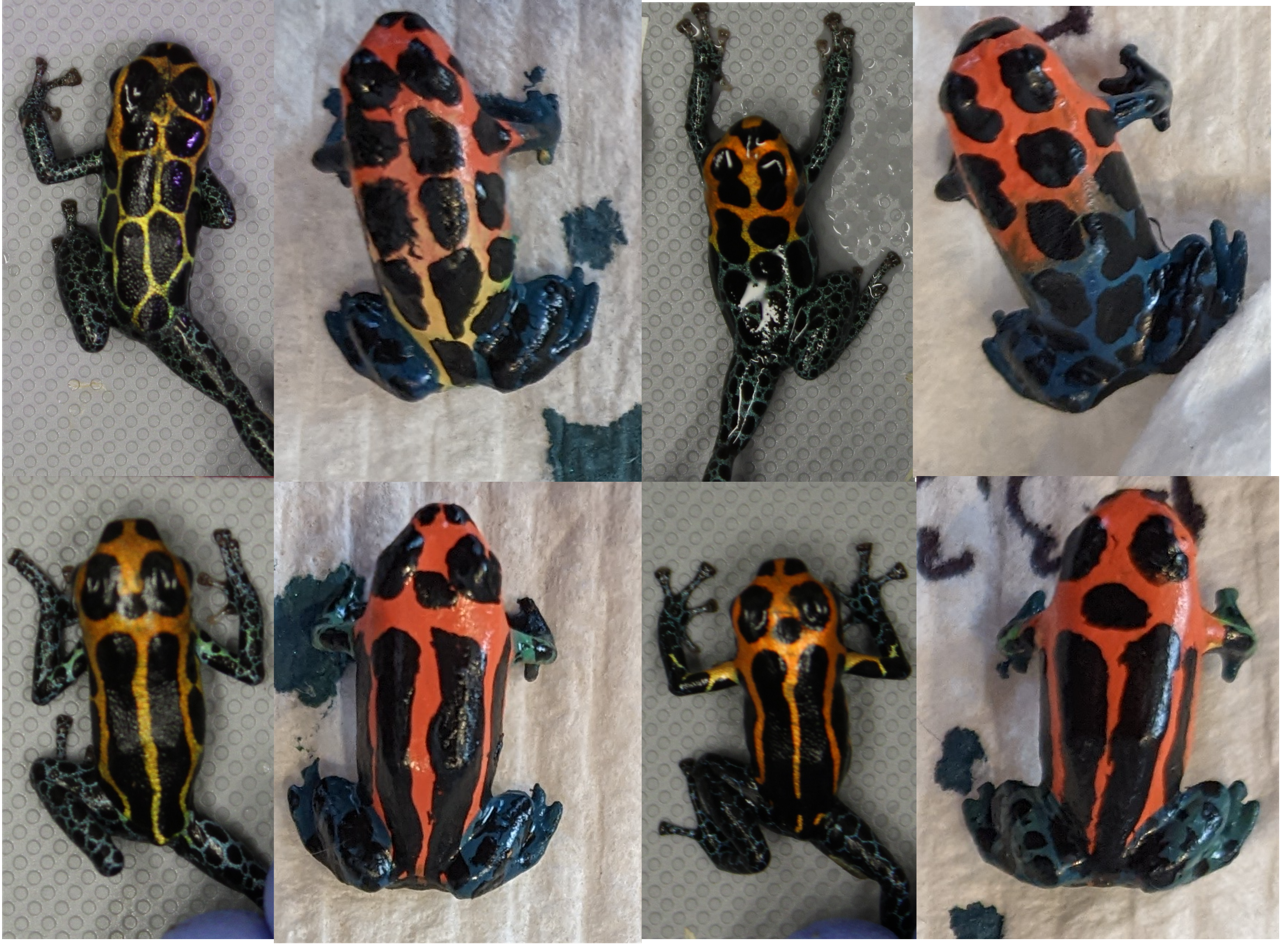

Figure S1: Dorsal views of individual male *R. imitator* (left) alongside of corresponding painted dummy frogs used as visual cues in mate removal trials (right).

B. *Acoustic Stimuli*

To generate acoustic stimuli for mate removal trials, calls were extracted from raw recordings of males made in our laboratory (Moss *et al.*, 2023). Male *R. imitator* produce distinct vocalizations for advertisement/territory defense, courtship, and parental care (Moss *et al.*, 2023). We previously showed that advertisement calls contain the most individual identity information (Moss *et al.*, 2023). Also, unlike short-range courtship and egg feeding calls, advertisement calls may function even when females are out of visual range. Indeed, pair-bonded females appear able to discriminate their male partners from strangers using advertisement calls alone (Podraza *et al.*, 2024). Thus, we used playback of advertisement calls to simulate individually distinctive soundscapes during mate removal trials.

Males were recorded in the laboratory following previously established protocols (Moss *et al.*, 2023; Podraza *et al.*, 2024). Briefly, subjects were recorded during daytime hours using a digital audio recorder (H4n Pro, Zoom, Tokyo, Japan) and shotgun microphone (K6/ME66, Sennheiser, Wennebostel, Germany) positioned atop the acoustically transparent screen mesh lid of the focal terrarium (<0.5 m from subjects). By cross-validating with video recordings, we have found these methods to be effective in capturing and isolating calls of focal individuals (Moss *et al.*, 2023)*.* Calls were recorded in WAV format at a sampling rate of 48 kHZ/16 bit. Based on preliminary analyses of recordings, we determined that males typically advertise in bouts of 2–80 calls and that as eggs approach hatching these bouts occur more frequently (i.e., ~every 15 minutes; see Fig. 1 of main text). To faithfully replicate this behavior and ensure that females with their mates removed experienced relatively constant acoustic stimulation from the time of male removal through the window for tadpole transport, whole bouts of advertisement calls, rather than singular advertisement calls, were used for playback.

Playback files were generated in the opensource audio editing software Audacity v.3.0.4 (https://www.audacityteam.org/). Bouts were identified as strings of advertisement calls, each separated by <1 min. After excising bouts from raw recordings, audio in between advertisement calls was manually silenced and Audacity’s ‘Noise Reduction’ tool was applied. Frequencies below 2,000 Hz and above 20,000 Hz were filtered using the ‘High Pass Filter’ and ‘Low Pass Filter’ tools, respectively. Once filtered, spacing was added between bouts such that the first call of the preceding bout occurred 15 minutes before the first call of the subsequent bout. In total, each playback file was 2 hours long and consisted of nine uninterrupted bouts.

C. *Scoring male call rates from trial video*

To quantify male call rates from video recordings of trials, we used the software, Wondershare Filmora (Wondershare Technology Group Co., Ltd, Vancouver, Canada) to splice around the time period of interest using the video as a guide and then exported the detached audio from this period. Detached audio was imported into Audacity v.3.0.4 (<https://www.audacityteam.org/>) for filtering and call quantification. We began by manually removing sections of audio afflicted by environmental noise indicating the presence of frog caretakers (i.e., doors opening and closing, noises generated during frog feeding, etc.). Frogs typically cease calling when disturbed, and therefore removing these periods of interruption effectively restricted the analysis to periods ‘suitable’ for calling. This process resulted in an average of 7.71 ± 3.58 hours of suitable time analyzed per trial (pre-hatching: 9.79 ± 2.23; hatch day: 7.11 ± 2.58; transport: 6.22 ± 4.70; range = 1.54–13.18). Additional filtering included application of the ‘Noise Reduction’, ‘High Pass Filter’ (<2000 Hz with 24 dB roll-off) and ‘Low Pass Filter’ (>20,000 Hz with 24 dB roll-off) tools. To facilitate visual detection of low amplitude calls in spectrograms, call quantification was carried out in Multiview mode. Calls were counted in 30-minute intervals and summed across the duration of the track. Focal calls were distinguished from background calls (i.e., originating from neighboring tanks) based primarily on the resolution of low frequency bands in the spectrograms and secondarily on sound (i.e., background calls contain little perceptible vibration or ‘vocal fry’).

D. *Transcriptome Assembly, Annotation, and Quantification*

After trimming adapter sequences, sequencing errors were corrected with RCorrector v1.0.7 with default parameters (Song & Florea, 2015). Corrected, paired-end reads were then supplied to Trinity v2.10.0 for *de novo* transcriptome assembly (Grabherr *et al.*, 2011). Following assembly, additional filtering steps included the clustering of redundant isoforms with CD-HIT v4.8.1 (Li & Godzik, 2006) and removal of short contigs (<250bp) with BBmap v39.01 (Bushnell, 2014). Finally, contaminant sequences (i.e., belonging to microorganisms and invertebrates) were removed by BLASTing contigs against the SwissProt database using a custom perl script (Alvarez-Buylla *et al.*, 2022). These filtering steps reduced the size of the assembly from an initial 2,238,044 contigs to a final 1,209,384 contigs. Final assembly completeness was evaluated with BUSCO v5.0.0 and estimated at 93.5% (39.2% duplicated; 1.4% fragmented; 5.1% missing).

Initial annotation of the assembly was performed using the Trinotate annotation pipeline (Bryant *et al.*, 2017), which identifies candidate coding regions in transcripts with Transdecoder v 5.5.0 (<https://github.com/TransDecoder/TransDecoder>) and annotates them, drawing from multiple databases. This process identified BLAST hits for 108,037 contigs. Additional annotation was performed using BLAST2GO by searching unannotated sequences of interest (i.e., transcripts differentially expressed between behavioral groups) against NCBI’s non-redundant vertebrate nucleotide database with an e-value cutoff of 1.0E-3. This brought the total number of annotated contigs to 108,960. Transcript quantification was performed using Salmon v 1.10.1 (Patro *et al.*, 2017) and alignment rates of each sample were summarized with MultiQC v1.14 (Ewels *et al.*, 2016).

*Supplementary Data Figures*

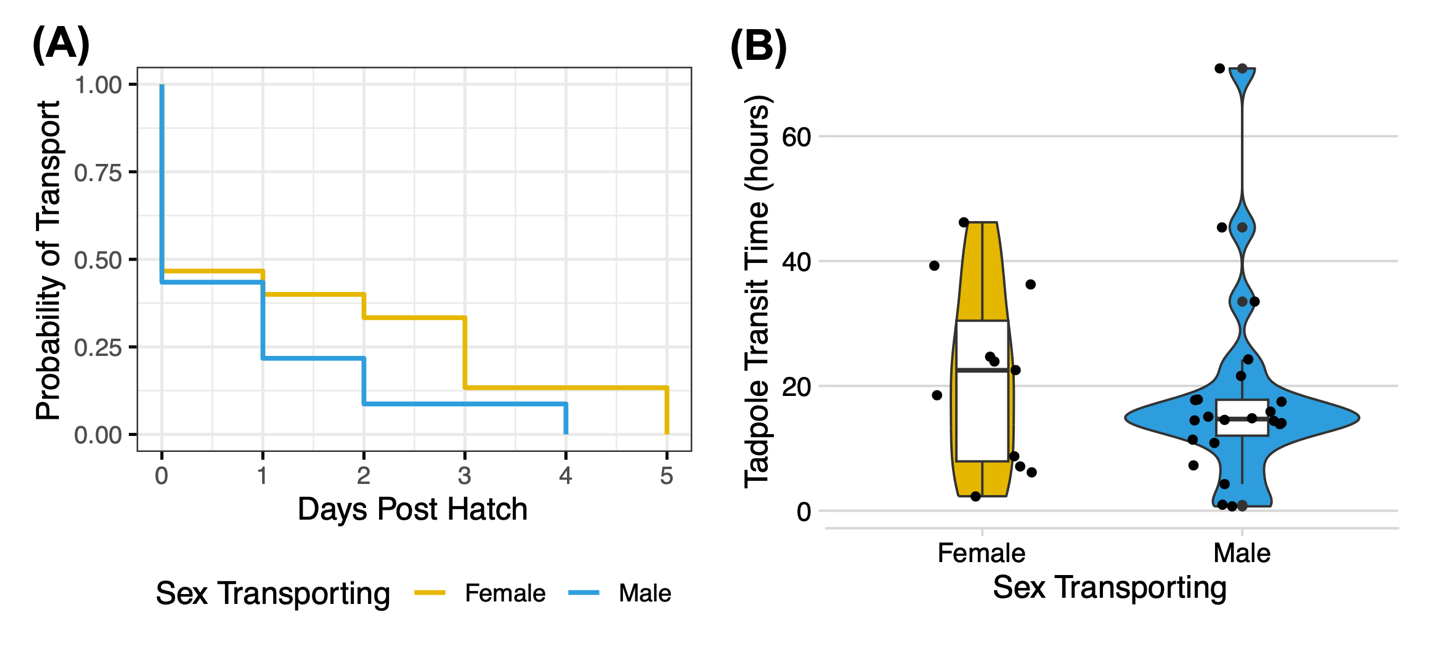

**Figure S2:** Tadpole transport behaviors compared between sexes. (A) Male and female survival curves depicting latency (post-hatching) to transport tadpoles, in days. (B) Male and female tadpole transit times, from pickup to deposition in a pool, in hours. There were no significant differences between the sexes in these metrics (Latency to transport: Likelihood ratio test=1.61, *P*=0.2; Transit Time: Χ^2^=0.2879; *P*=0.592).

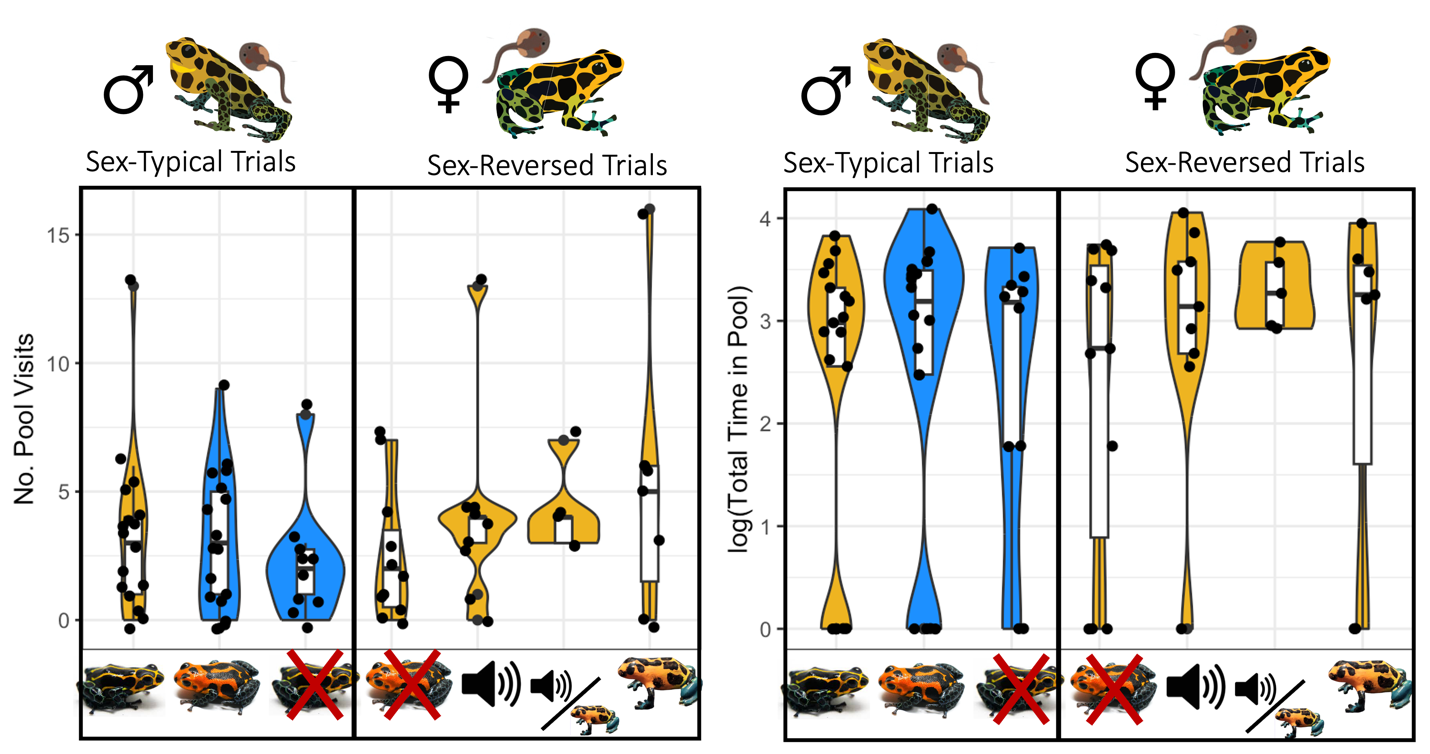

**Figure S3:** Pool visitation behavior on the last full day (12 hours) before the day of tadpole hatching, across trial types. Number of pool visits is shown on the left, and total time spent in pools is shown on the right. There were no significant differences between sexes or trial types in either pool visitation rates (Sex: Χ^2^=1.012, *P*=0.314; Trial Type: Χ^2^=4.364, *P*=0.359) or durations (Sex: Χ^2^=0.20, *P*=0.655; Trial Type: Χ^2^=0.526, *P*=0.261).

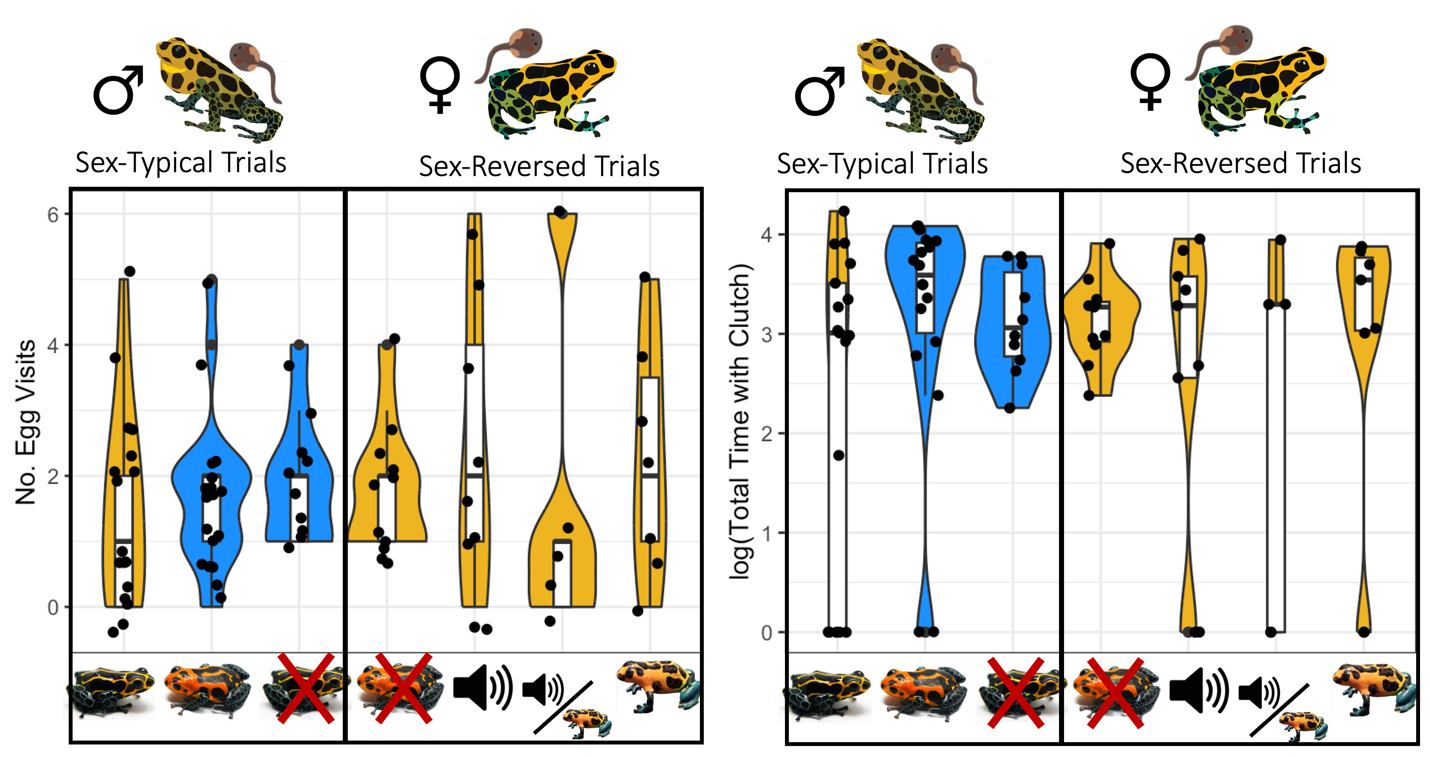

**Figure S4:** Egg visitation behavior on the last full day (12 hours) before the day of tadpole hatching, across trial types. Number of egg visits is shown on the left, and total time spent with eggs is shown on the right. There were no significant differences between sexes or trial types in either egg visitation rates (Sex: Χ^2^=0.009, *P*=0.924; Trial Type: Χ^2^ = 1.789, *P*=0.775) or durations (Sex: Χ^2^=2.813, *P*=0.094; Trial Type: Χ^2^=3.291, *P*=0.510).

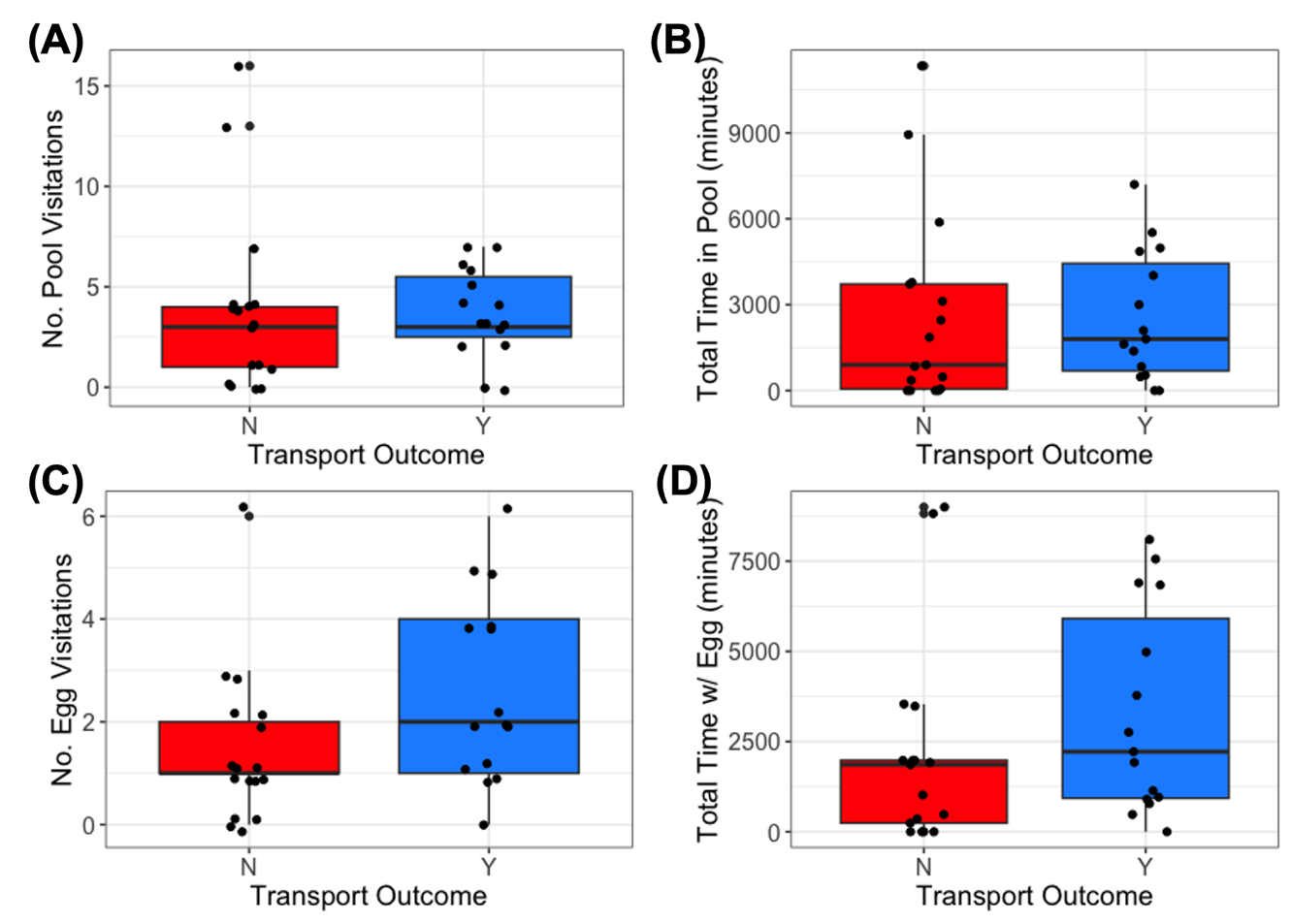

**Figure S5:** Distribution of prehatching parental behaviors on the last full day before egg hatching visualized in relation to the outcomes of sex-reversed trials (in terms of tadpole transport success by females): (A) Number of pool visitations (Χ^2^=0.225, *P*=0.635); (B) Total time in pools (Χ^2^=0.379, *P*=0.538); (C) Number of egg visitations (Χ^2^=2.811, *P*=0.094); (D) Total time with egg (Χ^2^=2.00, *P*=0.156).

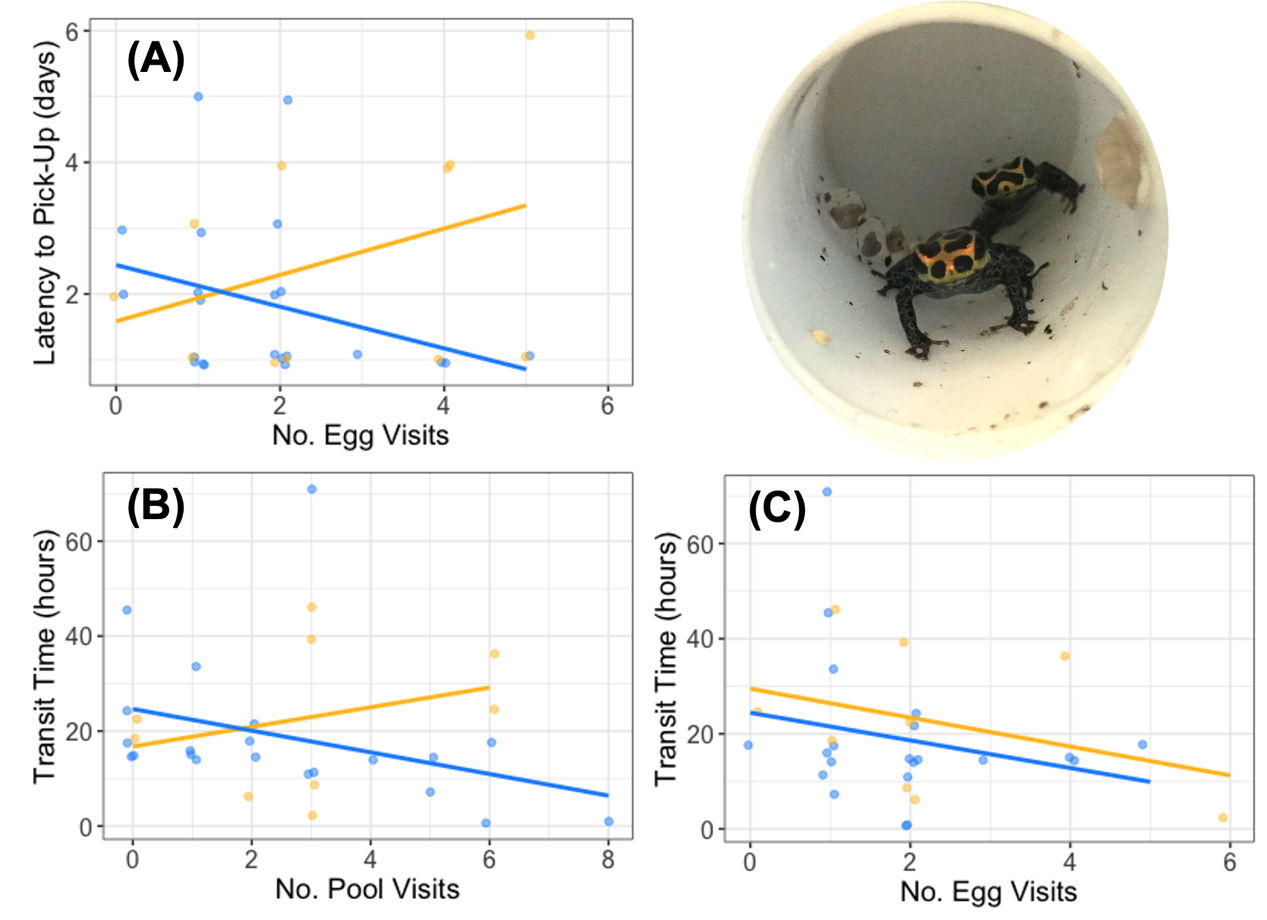

**Figure S6:** Significant regressions of post-hatching behaviors on pre-hatching behaviors on the last full day before egg hatching: (A) Latency to initiate transport (in days) regressed on number of egg visitations (Χ^2^=1.45; *P*=0.227) showed a marginally significant interaction with sex (Χ^2^=3.23; *P*=0.072); (B) Transit time (in hours) was significantly negatively associated with number of pool visitations (Χ^2^=4.668; *P*=0.031) but showed no sex=specific effects (Χ^2^=0.515; *P*=0.473); (C) Transit time (in hours) was significantly negatively associated with number of egg visitations (Χ^2^=15.943; *P*<0.0001) and this effect was more pronounced in females ((Χ^2^=8.014; *P*=0.005). For each plot, gold denotes female observations and blue denotes male observations.

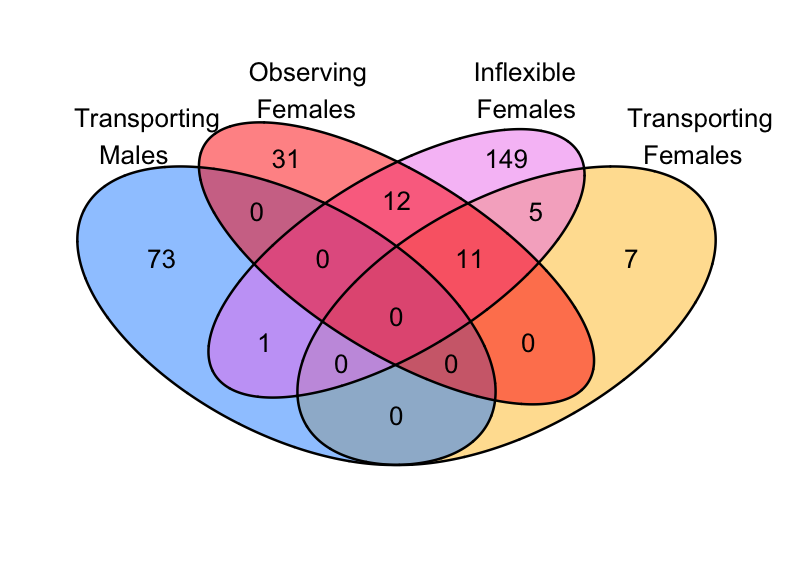

**Figure S7:** Differential brain gene expression across behavioral groups. Genes differentially expressed in pairwise comparisons to sex-matched egg care are shown in a Venn diagram with the number of distinct differentially expressed genes on edges and the number of shared differentially expressed genes in overlaps.

**Table S1:** Summary of individual information for *R. imitator* from behavioral experiment that underwent mate removal trials.

| **Frog ID** | **Morph** | **Source** | **Partner ID** | **Age at first trial (in days)** | **Experience at first Trial (in days since first lab clutch)** | **Experience at first Trial (in no. lab clutches)** | **Transported in Mate Removal?** |
| --- | --- | --- | --- | --- | --- | --- | --- |
| **Males** | | | | | | | |
| Ri.0172 | hybrid | Transferred as breeder | Ri.0173 | >167 | >43 | >5 | Y |
| Ri.0177 | hybrid | Transferred as breeder | Ri.0176 | >206 | >114 | >8 | Y |
| Ri.0181 | hybrid | Transferred as breeder | Ri.0205 | >315 | >147 | >13 | Y |
| Ri.0187 | hybrid | Transferred as breeder | Ri.1754 | >179 | >174 | >22 | Y |
| Ri.0188 | hybrid | Transferred as breeder | Ri.0174 | >550 | >288 | >22 | Y |
| Ri.0298 | green | Purchased as virgin (<6 mo) | Ri.0180 | 433 | 277 | 5 | Y |
| Ri.0300 | green | Purchased as virgin (<6 mo) | Ri.0182 | 97 | 56 | 3 | Y |
| Ri.0442 | veradero | Purchased as virgin (<6 mo) | Ri.1841 | 210 | 15 | 2 | Y |
| Ri.0464 | hybrid | Lab born | Ri.1188 | 368 | 43 | 5 | Y |
| Ri.0981 | veradero | Purchased as virgin (<6 mo) | Ri.1482 | 273 | 123 | 7 | Y |
| Ri.1562 | hybrid | Lab born | Ri.0540 | 179 | 11 | 3 | Y |
| **Females** | | | | | | | |
| Ri.0173 | hybrid | Transferred as breeder | Ri.0172 | >167 | >43 | >5 | Y |
| Ri.0176 | hybrid | Transferred as breeder | Ri.0185 | >206 | >114 | >8 | N |
| Ri.0178 | hybrid | Transferred as breeder | Ri.1761 | >416 | >233 | >11 | N |
| Ri.0182 | hybrid | Transferred as breeder | Ri.0300 | >168 | >49 | >7 | Y |
| Ri.0184 | hybrid | Transferred as breeder | Ri.1643 | >169 | >152 | >9 | Y |
| Ri.0186 | hybrid | Transferred as breeder | Ri.0187 | >179 | >174 | >22 | Y |
| Ri.0205 | hybrid | Transferred as breeder | Ri.0181 | >286 | >55 | 8 | N |
| Ri.0441 | veradero | Purchased as virgin (<6 mo) | Ri.0442 | 210 | 15 | 2 | Y |
| Ri.0516 | hybrid | Lab born | Ri.0981 | 503 | 80 | 3 | N |
| Ri.0525 | hybrid | Lab born | Ri.0975 | 523 | 245 | 8 | Y |
| Ri.0543 | hybrid | Lab born | Ri.0298 | 320 | 111 | 3 | N |
| Ri.0973 | veradero | Purchased as virgin (<6 mo) | Ri.1680 | 131 | 17 | 2 | Y |
| Ri.0982 | veradero | Purchased as virgin (<6 mo) | Ri.0464 | 119 | 43 | 5 | Y |
